# Skeletal muscle mitochondrial dysfunction in mice is linked to bone loss via the bone marrow immune microenvironment

**DOI:** 10.1101/2020.12.09.417147

**Authors:** Jingwen Tian, Ji Sun Moon, Ha Thi Nga, Hyo Kyun Chung, Ho Yeop Lee, Jung Tae Kim, Joon Young Chang, Seul Gi Kang, Dongryeol Ryu, Xiangguo Che, Je-Yong Choi, Masayuki Tsukasaki, Takayoshi Sasako, Sang-Hee Lee, Minho Shong, Hyon-Seung Yi

**Author notes:** Corresponding author: Hyon-Seung Yi, Research Center for Endocrine and Metabolic Diseases, Chungnam National University School of Medicine, Daejeon 35015, Korea.; **E-mail**.

## Abstract

Mitochondrial oxidative phosphorylation (OxPhos) is a critical regulator of skeletal muscle mass and function. Although muscle atrophy due to mitochondrial dysfunction is closely associated with bone loss caused by reduction of mechanical loading, questions remain about the biological characteristics in the relationship between muscle and bone. Here, we have shown that muscle atrophy caused by skeletal muscle-specific *Crif1* knockout (MKO) modulates the bone marrow inflammatory response, leading to bone loss. Transcriptome analysis of the extensor digitorum longus revealed that local mitochondrial stress increased serum levels of fibroblast growth factor 21 (FGF21) in mice. However, we have shown by *Fgf21* knockout in MKO mice that FGF21 is dispensable for muscle atrophy-mediated bone loss. RNA sequencing in MKO mice indicated that mitochondrial stress response in skeletal muscles induces an inflammatory response and adipogenesis in the bone marrow. We also found, using transcriptomic analysis of bone marrow, that the CXCL12–CXCR4 axis is important for T-cell homing to the bone marrow, which is an immunological mediator of muscle-bone communication. CXCR4 antagonism attenuated bone marrow inflammation and bone loss in MKO mice. Together, these data highlight the role that muscle mitochondrial dysfunction plays in triggering bone marrow inflammation via the CXCL12–CXCR4 signaling axis, which is critical for inducing bone loss.

## Introduction

Mitochondria quality control pathways regulating mitochondrial morphology/function have been implicated in the homeostatic control of muscle mass ^1^. Gradual loss of muscle mass and strength results from an inflammatory state brought about by cytokines, apoptosis, alteration in mitochondrial respiration, and oxidative stress ^2^. Decrease or dysfunction of skeletal muscle mitochondria is involved in loss of muscle mass and muscle strength in humans ^3^. Progressive loss of muscle mass and function has also been implicated as a critical risk factor for osteoporosis through reduction of bone strength caused by decreasing mechanical loading on the skeleton ^4,5^. Moreover, mitochondrial DNA-mutator mice showed premature aging-related phenotypes including muscle dysfunction and osteoporosis ^6^. However, little information is available on the mechanical link between mitochondrial dysfunction-mediated loss of muscle mass and strength, and bone loss.

Owing to their proximities, bone marrow (BM) immune cells contribute to the regulation of osteoblasts and osteoclasts, leading to modulation of bone remodeling. For example, ovariectomy induces T-cell activation and proliferation in BM, which promotes osteoclastogenesis ^7^ due to increases in the populations of CD4^+^ and CD8^+^ T-cells, and in T-cell derived tumor necrosis factor alpha (TNF-α). Additionally, ovariectomy-induced bone loss does not occur in mice lacking T-cells or lacking the T-cell receptor CD40 ligand ^8,9^. Germ-free mice have a smaller population of CD4^+^ T-cells and reduced levels of proinflammatory cytokines in the BM in combination with increased bone mass ^10^. Moreover, BM CD4^+^ T-cells producing interleukin (IL)-17 and TNF-α, but not interferon (IFN)-γ, activate bone resorption by inducing osteoclast differentiation in animal models of inflammatory bowel disease ^11^. However, although previous investigations revealed several factors promoting BM inflammatory-mediated bone loss, the role of mitochondrial function in skeletal muscle on the BM immune microenvironment and bone mass remains to be elucidated.

CR6-interacting factor 1 (CRIF1), as a critical mitoribosomal protein, is essential for the translation of mitochondrial oxidative phosphorylation (OxPhos) subunits and their insertion in the mitochondrial inner membrane ^12^. Deficiency of CRIF1 produces impaired formation of the OxPhos complex, leading to reduction of mitochondrial respiration. Consequently, both beta cell- and adipocyte-specific *Crif1* knockout mice show mitochondrial OxPhos dysfunction-associated metabolic phenotypes such as glucose intolerance and adipose inflammation ^13,14^. On the other hand, we have shown that OxPhos dysfunction in skeletal muscle-specific *Crif1* knockout (MKO) mice is associated with improved glucose tolerance and insulin resistance ^15^. However, in that study, we found that MKO mice had markedly smaller gastrocnemius and extensor digitorum longus (EDL) muscles compared with the controls ^15^. Thus, we hypothesized that MKO mice could be used as models for studying the mechanisms underlying muscular mitochondrial dysfunction-induced bone loss.

Herein, we investigated whether skeletal muscle dysfunction caused by muscle-specific OxPhos deficiency changes the population of inflammatory T-cells and the levels of proinflammatory cytokines in the BM, thereby resulting in bone loss. We also examined the role of key mitokines, produced in response to mitochondrial stress, in the regulation of bone mass and mineral density. In addition, the present study identifies the mechanism underlying deterioration in bone mass and quality caused by loss of muscular mitochondrial function and, consequently, a potential therapeutic target.

## Results

### Crif1 deficiency induces defective mitochondrial OxPhos function and stress response in skeletal muscles

To assess the effect of skeletal muscle-specific *Crif1* deficiency on the mitochondrial function and physical performance of the mice, we generated MKO mice by crossing *Mlc1f-cre* mice with *Crif1*^*flox/flox*^ mice (Supplementary Fig 1a). CRIF1 protein expression was markedly lower in the EDL, gastrocnemius, and soleus of the MKO mice than in the wild-type (WT) controls (Fig 1a,b, and Supplementary Fig 1b,c). Consistent with the reduction in CRIF1 expression, the EDL and gastrocnemius in MKO mice showed significantly lower expression of mitochondrial OxPhos complex subunits, including complex II (SDHA), III (UQCRC2), and V (ATP5A), indicating that deficiency of *Crif1* induces abnormal mitochondrial proliferation and function in skeletal muscles (Fig 1a,b, and Supplementary Fig 1b,c). Moreover, blue native polyacrylamide gel electrophoresis (BN-PAGE) analysis revealed a decrease in assembly of complex I, III, and V in MKO mouse EDL and gastrocnemius compared with the WT (Fig 1c). MKO mice also exhibited lower succinate dehydrogenase (SDH) activity in the EDL, gastrocnemius, and soleus compared with the WT (Fig 1d,e, and Supplementary Fig 1d). Electron microscopy also revealed accumulation of abnormal, swollen mitochondria with disrupted cristae in gastrocnemius of MKO mice (Supplementary Fig 1d). In addition, EDL from MKO mice expressed higher levels of mitochondrial stress response-related genes, such as *Lonp1*, *Clpp*, *Hspd1*, and *Atf4* (Fig 1f).

**Figure 1.**
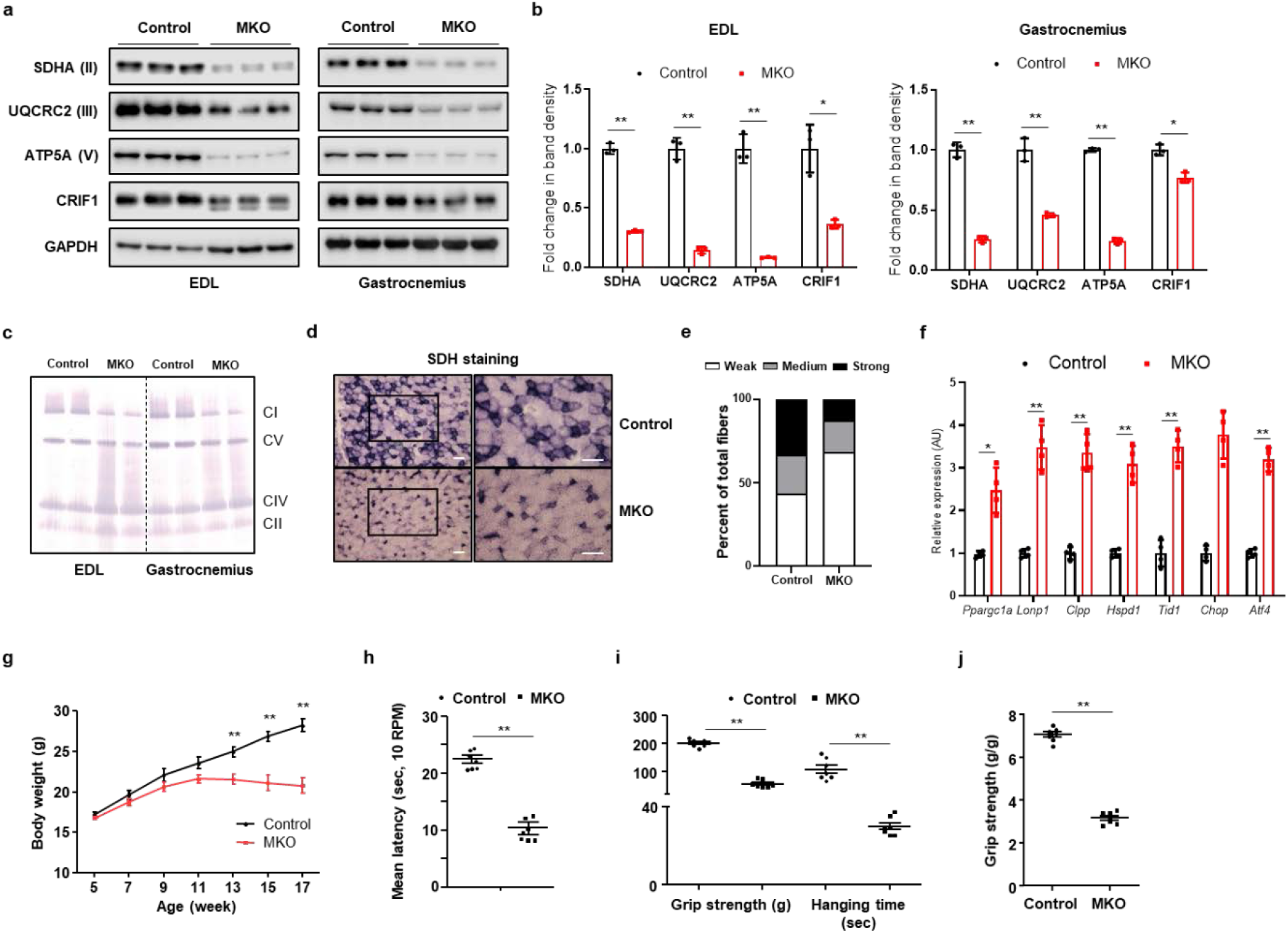
MKO mice show impairment in OxPhos and deterioration in physical performance. (a, b) Representative western blots and band density measurements for OxPhos complex subunits and CRIF1 in the EDL and gastrocnemius of chow-fed control and MKO mice at 14 weeks of age (*n* = 3). (c) Representative blots showing BN-PAGE of the assembled OxPhos complex in EDL and gastrocnemius from chow-fed control and MKO mice at 14 weeks of age. (d) Transverse EDL sections were histochemically stained for SDH to identify oxidative muscle fibers at 14 weeks of age. Scale bar, 100 μm. (e) Quantification of unstained fibers in EDL of controls and MKO mice at 14 weeks of age (each *n* = 5). (f) Relative mRNA expression of genes related to mitochondrial stress response from EDL in 14-week-old control and MKO mice (each *n* = 4). (g) Body weight evolution of control and MKO mice fed a chow diet for 8 weeks (*n* = 10 per group). (h) Latency to fall in rotarod test at 10 rpm (each *n* = 7). (i) Forelimb grip strength and time to fall in the wire hanging assay for control and MKO mice (each *n* = 10). (j) Grip strength normalized to body weight of control and MKO mice (each *n* = 10). Data are expressed as mean ± SEM. Statistical significance was analyzed by unpaired t-tests. *, *P* < 0.05 and **, *P* < 0.01 compared with the indicated group.

Next, we characterized the physical performance of MKO mice on a normal chow diet. MKO mice showed a significant decrease in body mass relative to controls after 11 weeks of age (Fig 1g). Motor coordination was assessed using rotarod apparatus. At fixed rotarod speed (10 rpm), a significant difference between control and MKO mice was observed in latency to fall (Fig 1h). Furthermore, unlike the controls, MKO mice showed a decline in grip strength and a higher drop rate in the wire hanging test at 13 weeks of age (Fig 1i,j). Collectively, these findings indicate that *Crif1* deficiency not only adversely affects mitochondrial OxPhos function, but also reduces muscle strength and endurance in mice.

### Mitochondrial OxPhos dysfunction in muscle promotes bone loss in mice

Loss of muscle mass and function is closely associated with increased bone loss and fracture incidence ^16^. Thus, to elucidate the crosstalk between mitochondrial function of skeletal muscle and bone mass, at 14 weeks of age we investigated the bone mass and mineral density (BMD) of control and MKO mice on a normal chow diet. Analysis of femurs from euthanized mice by micro-computed tomography (micro-CT) revealed that cortical bone thickness and BMD were significantly decreased in MKO mice (Fig 2a and Supplementary Fig 2a). Femurs from MKO mice also showed a decrease in trabecular number, trabecular thickness (Tb.Th.), and trabecular bone volume (BV), but exhibited an increase in trabecular spacing (Tb.Sp.) compared with those from the controls (Fig 2b,c). Quantification and statistical analysis of the bone phenotypes characterized by micro-CT are shown in Figure 2C.

**Figure 2.**
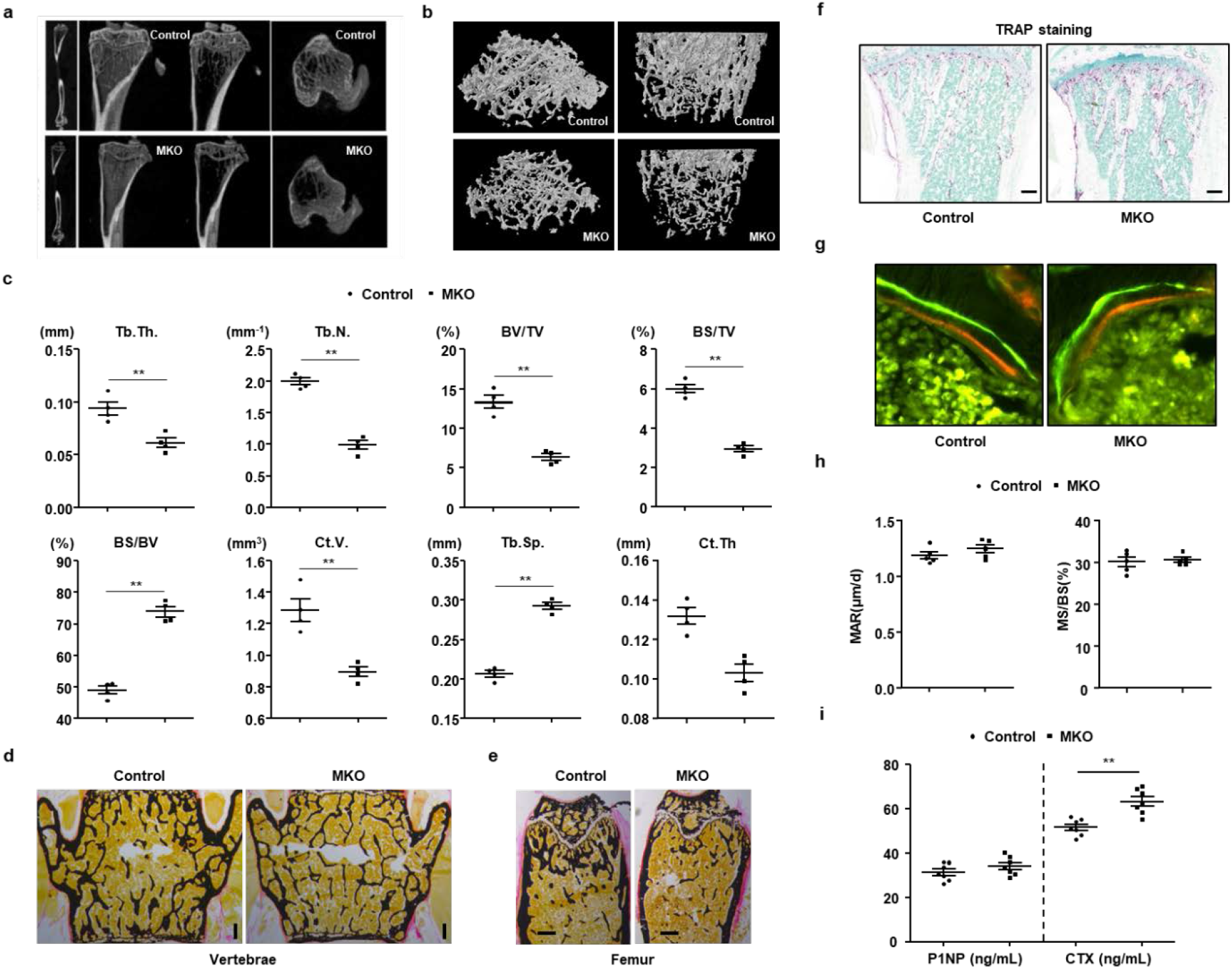
Mitochondrial stress response in skeletal muscle promotes bone loss. (a,b) Representative images of micro-CT of cortical and trabecular regions in the distal femur from control and MKO mice at 14 weeks of age. (c) Measurement of Tb.Th., trabecular number (Tb.N.), BV/TV, BS/TV, BS/BV, cortical volume (Ct.V.), Tb.Sp., and total bone volume (TBV) using micro-CT analysis. (d,e) Von Kossa staining of undecalcified sections of vertebrae and femurs of control and MKO mice at 14 weeks of age. Scale bars, 250 μm. (f) TRAP staining to reveal osteoclasts. Scale bars: 100 μm. (g) Histomorphometric analysis of calcein/alizarin red double-stained sections was conducted to quantify bone formation in vertebrae (*n* = 5/group). Green indicates calcein, and red indicates alizarin red. (h) Measurement of endocortical MAR and MS/BS (%) of vertebrae in control and MKO mice. (i) Serum levels of P1NP and CTX in 14-week-old control and MKO mice (each *n* = 7). Data are expressed as mean ± SEM. Statistical significance was analyzed by unpaired t-tests. *, *P* < 0.05 and **, *P* < 0.01 compared with the indicated group.

Bone histomorphometry verified that MKO mice developed a low bone-mass phenotype in cortical and trabecular bone, as shown by von Kossa staining (Fig 2d,e, and Supplementary Fig 2b,c), but they showed a high number of osteoclasts per bone surface, as indicated by tartrate-resistant acid phosphatase (TRAP)-stained sections of tibia (Fig 2f and Supplementary Fig 2d). Next, to quantify the extent of increased osteoblast activity, calcein and alizarin red were sequentially given to control and MKO mice 3 and 8 days prior to sacrifice. Analysis of undecalcified, unstained sections showed no notable difference in vertebral bone formation rates between control and MKO mice (Fig 2g,h). Additionally, measurement of the serum level of procollagen type I N-terminal propeptide (P1NP), a marker for bone formation, and of C-terminal telopeptide of collagen (CTX), a marker for bone resorption, indicated less bone formation and higher bone resorption in MKO mice compared with the controls (Fig 2i). To exclude the effect of other endocrine hormones on bone loss, we checked the serum levels of parathyroid hormone, testosterone, thyroxine, and triiodothyronine in the control and MKO mice at 14 weeks of age. There was no significant difference in the serum level of each of these hormones between the control and MKO mice (Supplemental Fig 3a-d). Taken together, these data provide direct evidence that lower mitochondrial OxPhos-mediated muscle dysfunction induces bone loss via activation of osteoclasts in mice.

### Bone loss in MKO mice is independent of FGF21 action

The phenotypic changes observed so far suggested that MKO mice express circulating factors involved in signaling from muscle to bone. Therefore, to study the effects of muscular mitochondrial dysfunction-induced secretory factors on bone loss in MKO mice, we performed RNA sequencing of EDL transcripts from control and MKO mice, focusing on genes encoding secreted proteins. A large number of transcripts were altered in the EDL from MKO mice (Fig 3a,b). In particular, we found that the expression of *Fgf21*, which is known as a potent regulator of skeletal homeostasis, was much higher in the EDL of MKO mice than in that of the controls (Fig 3b-d). As shown in Figure 3E, the serum levels of FGF21 were also markedly elevated in the MKO mice.

**Figure 3.**
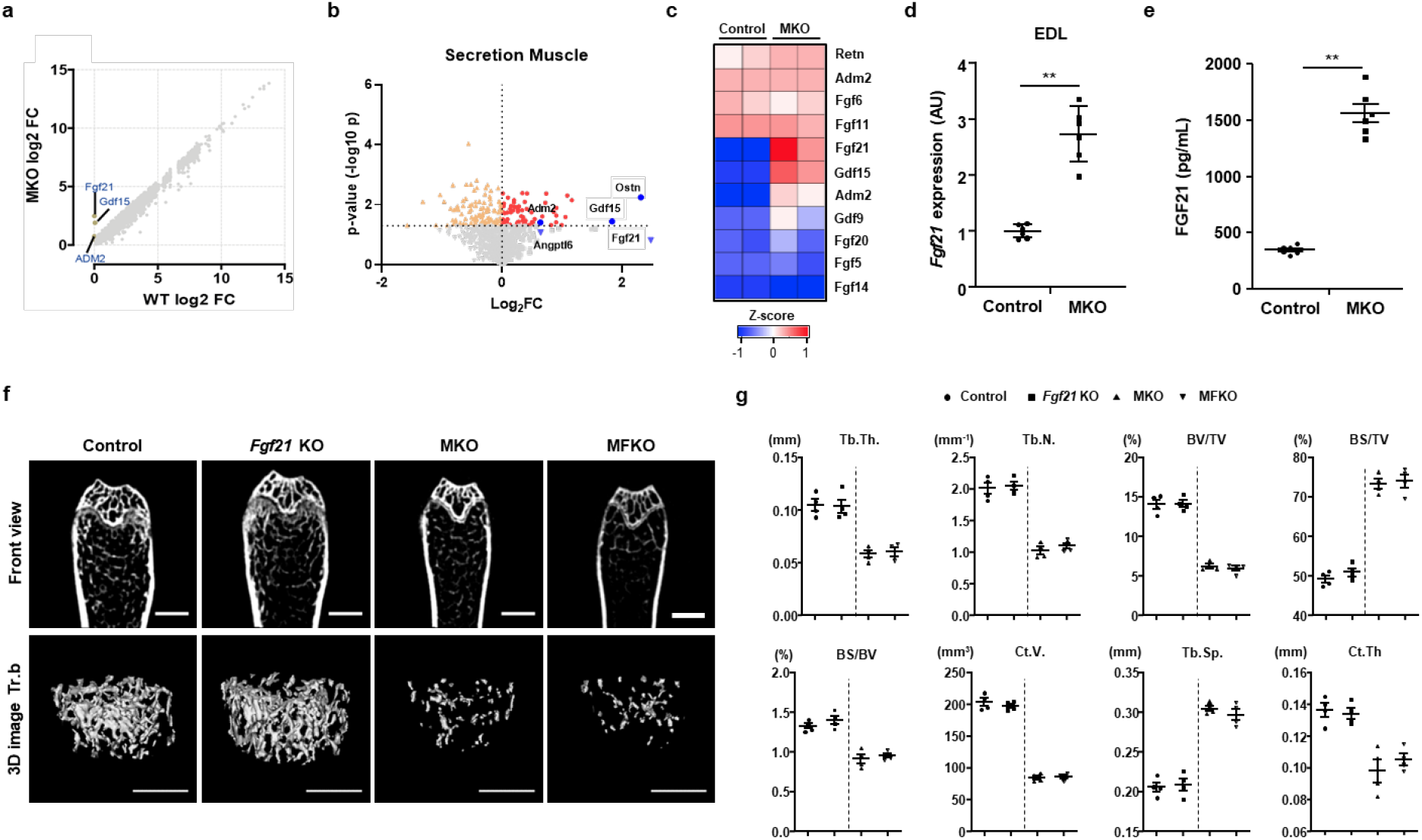
Bone loss is independent of FGF21 production caused by mitochondrial stress in skeletal muscle of MKO mice. (a)Scatterplots of RNA sequencing data, displaying transcript levels in EDL of control (x-axis) and MKO (y-axis) mice at 10 weeks of age. The text indicates mitokines showing much higher fold-change in MKO mice among the secreted proteins. (b,c) Volcano plot and heat map showing upregulated genes in EDL from normal chow diet-fed control and MKO mice at 10 weeks of age. (d) Relative expression of mRNA encoding *Fgf21* in EDL from 14-week-old control (*n* = 6) and MKO (*n* = 6) mice. (e) Serum levels of FGF21 in 14-week-old control (*n* = 6) and MKO (*n* = 6) mice. (f) Micro-CT images of the trabecular bone (Tr.b) near the distal femoral metaphyseal region from control (*n* = 4) and MKO (*n* = 4) mice at 14 week of age. Scale bar for front view and 3D image Tr.b, 1000 and 500 μm, respectively. (g) Measurement of Tb.Th., Tb.N., BV/TV, BS/TV, BS/BV, Ct.V., Tb.Sp., and TBV using micro-CT analysis. Data are expressed as mean ± SEM. Statistical significance was analyzed by unpaired t-tests. **, *P* < 0.01 compared with the indicated group.

Next, to investigate the role of FGF21 on the skeletal phenotype in MKO mice, we generated global *Fgf21* knockout mice and MKO mice with global *Fgf21* deletion (MFKO), both on a C57BL/6J background (Supplementary Fig 4a,b). Analysis of femurs by micro-CT revealed that cortical thickness, Tb.Th., and trabecular number were remarkably lower in MKO mice compared with the controls, but global *Fgf21* knockout did not have any effect on the bone phenotype of WT control or MKO mice at 14 weeks of age (Fig 3f,g). These findings suggest that a reduction of BMD in MKO mice is independent of FGF21 production caused by muscular mitochondrial OxPhos dysfunction.

### MKO mice exhibit inflammation response in BM

In the next set of experiments, we asked whether mitochondrial OxPhos dysfunction in muscle affects BM inflammation, which is an important factor for instigating bone resorption. To address this issue, we investigated the expression of proinflammatory cytokines in the BM cells from control and MKO mice. The expression of *Tnf*, *Rankl*, *Rorat*, and *Il17a* was significantly increased in the BM from MKO mice compared with the controls (Fig 4a). Seeking further evidence in support of the concept that mitochondrial OxPhos dysfunction in skeletal muscle induces inflammation in BM, we measured the populations of diverse T-cells in the BM from control and MKO mice using flow cytometry analysis. We found that at 14 weeks of age, mature CD3+ T-cells were markedly elevated in the BM from MKO mice compared with those from the controls (Fig 4b,c). Moreover, the population of TNF-α-producing CD4+ and CD8+ T-cells was significantly larger in the BM of MKO mice (Fig 4d,e). The populations of CD44+ TNF-α+ and CD44+IL-17A+ cells were also larger among the CD4+ and CD8+ T-cells of the BM of MKO mice (Fig 4d-h). However, there was no difference in the size of the population of BM CD4+CD25+Foxp3+ regulatory T-cells between control and MKO mice (Fig 4i). Furthermore, to exclude systemic inflammation underlying the bone loss in MKO mice, we measured TNF-α and IL-17A production in CD4+ T-cells from the spleen of control and MKO mice at 14 weeks of age. The expression of proinflammatory cytokines by CD4+ T-cells was not different in the spleens of control and MKO mice (Fig 4j-l). In addition, there was no notable difference in the serum level of TNF-α or IL-17A between control and MKO mice at 14 weeks of age (Supplementary Fig 5). Thus, these data revealed that muscular mitochondrial dysfunction-mediated bone loss is associated with BM inflammation rather than a systemic inflammatory response.

**Figure 4.**
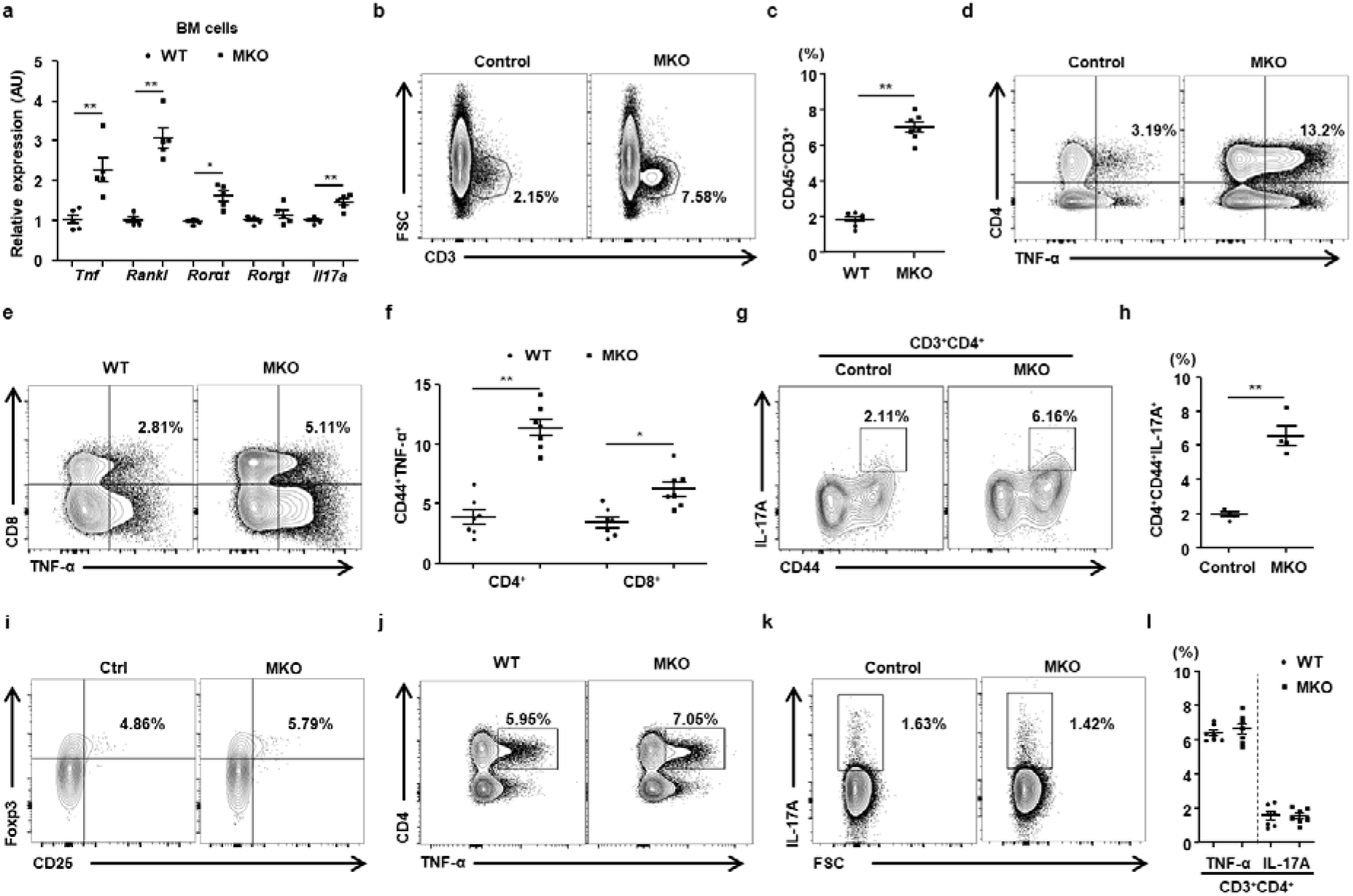
MKO mice exhibit local induction of TNF-α and IL-17 in BM without a systemic inflammatory response. (a) BM cells isolated from control (*n* = 5) and MKO mice (*n* = 5) at 14 weeks of age were subjected to real-time PCR analysis of osteoclastogenic genes. (b) Representative flow cytometry contour plots are presented for CD3 expression by BM cells in control (*n* = 7) and MKO (*n* = 7) mice at 14 weeks of age. (c) Statistical analysis of the population of CD3+ T-cells in BM cells in the two groups. (d-h) The number of TNF-α- or IL-17A-secreting cells in the populations of BM CD4+CD44+ and CD8+C44+ T-cells was compared between the two groups (each *n* = 7). (i) Population of regulatory T-cells (CD4+CD25+FOXP3+) in the BM of 14-week-old control (*n* = 7) and MKO (*n* = 7) mice. (j-l) TNF-α- or IL-17A-producing cells among CD4+ and CD8+ T-cells from the spleens of control (*n* = 7) and MKO (*n* = 7) mice at 14 weeks of age. Data are expressed as mean ± SEM. Statistical significance was analyzed by unpaired t-tests. *, *P* < 0.05 and **, *P* < 0.01 compared with the indicated group.

### MKO mice show adipogenesis and an inflammatory response in the BM

Based on the results from the flow cytometry analysis of the BM of control and MKO mice, we further investigated the transcripts of whole BM cells using RNA sequencing. Given that macroscopically the long bones dissected from MKO mice were more reddish than the controls (Fig 5a), we assumed that a more intense inflammatory response had occurred in the BM of MKO mice. After hematoxylin and eosin (H&E) staining, we also discovered a higher number of lipid droplets in the BM of MKO mice (Fig 5b). Thus, we conducted RNA sequencing to identify genes in BM cells differentially expressed between control and MKO mice at 14 weeks of age. Enrichment analysis with Network2Canvas revealed that genes associated with myopathy, osteoporosis, signaling in the immune system, T-cells, and IL-12 and IL-17 pathways were enriched in the BM cells from MKO mice (Fig 5c, and Supplementary Fig 6a,b). Moreover, gene set enrichment analysis (GSEA) indicated that the gene sets involved in adipogenesis or adipocyte maturation was enriched in the BM cells of MKO mice compared with the controls (Fig 5d,e). In addition, consistent with the long bones appearing more reddish in MKO mice, genes related to T-cell activation and co-stimulation were markedly increased in the BM cells of MKO mice, which may be closely associated with the BM inflammation and bone loss observed (Fig 5f,g). A volcano plot based on differential expression and statistical significance of the difference showed that, in addition to that of the adipogenesis and T-cell activation gene sets, expression of *Cxcl12* as well as of *Fabp4*, *Adipoq*, and *Cd3e* was also changed in the BM cells of MKO mice (Fig 5h). We also found that CXCL12 protein expression was higher in the PDGFRβ+VCAM-1+ stromal cells (CXCL12-rich cells) among BM cells from MKO mice using flow cytometry analysis (Fig 5i and Supplementary Fig 6c). Furthermore, we searched genes potentially associated with *Cxcl12* by applying Gene-Module Association Determination (G-MAD) to mouse expression data sets using GeneBridge tools ^17^. As a result, genes annotated in the inflammatory response and T-cell activation functional clusters were strongly enriched (Supplementary Fig 6d). These results implicate the distinct effects of muscular mitochondrial dysfunction on BM inflammation and bone loss in mice.

**Figure 5.**
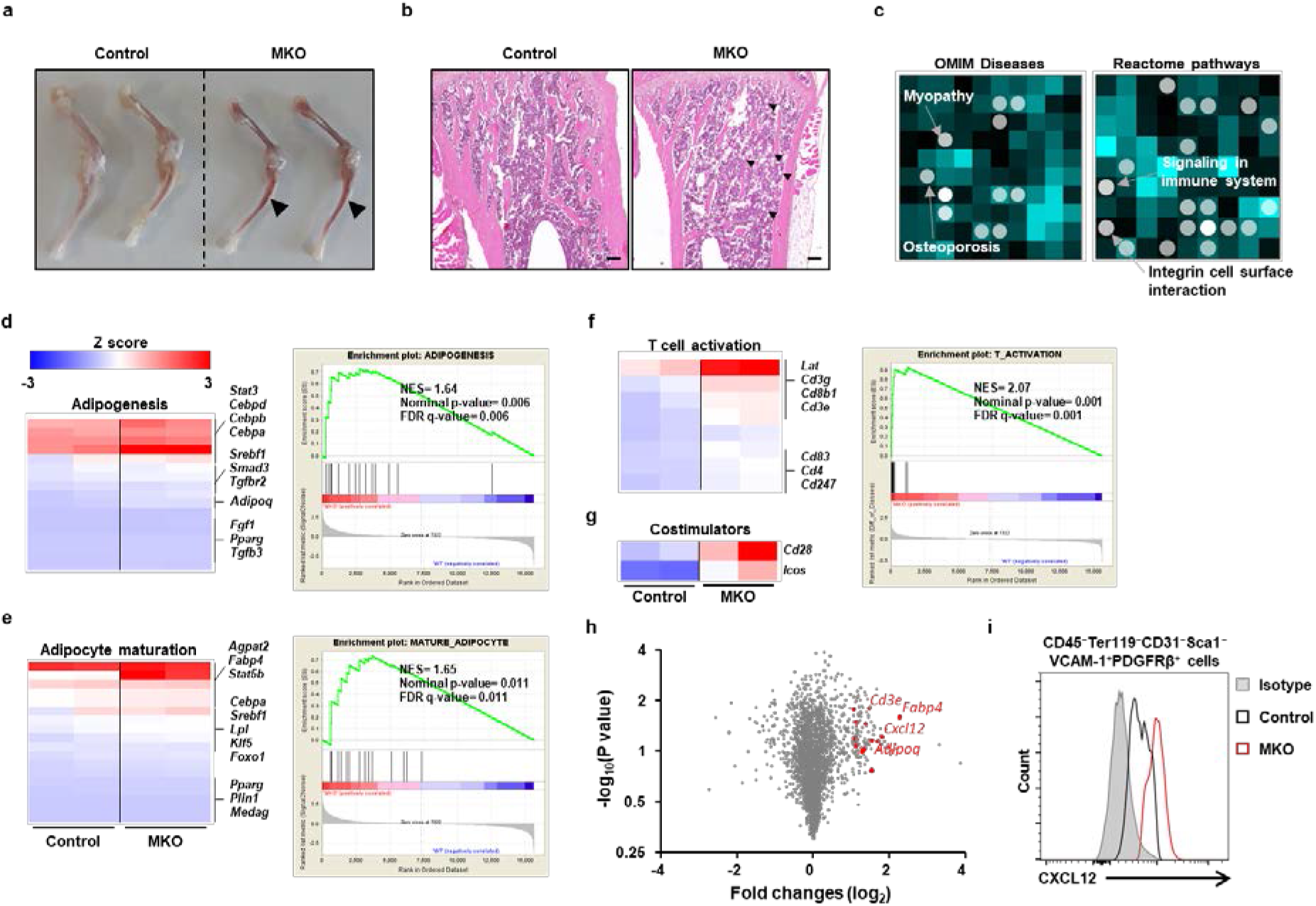
Transcriptome analysis shows adipogenesis and inflammatory response in the BM of MKO mice. (a) Femur and tibia of a 14-week-old control and MKO mouse. BM is more visible through the thin cortical bone of MKO mice. (b) A representative section of tibia from a 14‐week‐old control and MKO mouse stained with H&E. Thin femoral bone and adipocyte-rich BM (arrowhead) are visible in the MKO femur. Scale bars, 250 μm. (c) Using RNA sequencing data, genes that were significantly upregulated in the BM cells of control and MKO mice were analyzed for gene-list enrichment with gene set libraries created from level 4 of the MGI mouse phenotype ontology using Network2Canvas. (d-g) The diagram shows the results of gene set (Adipogenesis, Adipocyte maturation, T-cell activation, and Co-stimulators) enrichment analysis, including the enrichment scores. (h) Volcano plot based on the differential expression and significance of the differences in the data from RNA sequencing of BM cells of control and MKO mice at 14 weeks of age. Red dots represent genes associated with adipogenesis and T-cell activation with a *P*-value <0.05 and a fold-change >1 log2. (i) In the flow cytometry analysis, total BM cells were gated for a population negative for CD45, Ter119, CD31, and Sca1, and were positive for VCAM and PDGFRβ, the phenotype of CXCL12-abundant reticular cells. Intracellular levels of CXCL12 are shown as amount of protein in CAR cells in the BM of MKO mice relative to CAR cells in the BM of the controls at 14 weeks of age (*n* = 4 mice/genotype). Scale bars: 50 μm (a) and 10 μm (b). Data are expressed as mean ± SEM. Statistical significance was analyzed by unpaired t-tests. *, *P* < 0.05 and **, *P* < 0.01 compared with the indicated group.

### CXCR4 antagonism attenuates BM inflammation in MKO mice

CXCL12 is also strongly expressed in the reticular cells located adjacent to sinusoids in BM, which express adiponectin and are targetable with an adiponectin-Cre transgene ^18^. In addition, differentiated adipocytes induce CXCL12 secretion, which recruits proinflammatory macrophages into the adipose tissue of diet-induced obese mice ^19^. The chemokine CXCL12 binds primarily to CXCR4, leading to activation of intracellular signaling in multiple cell types including lymphocytes ^20^. Moreover, the CXCL12–CXCR4 axis is involved in many physiological and pathological processes ^20^. Thus, we asked whether migration of CXCR4+ immune cells to the MKO mouse BM contributes to the inflammatory response and proinflammatory cytokine production therein. As shown in Figure 6A, G-MAD revealed that genes annotated in the inflammatory response and T-cell activation functional clusters were strongly enriched. Next, to reveal the function of the CXCL12–CXCR4 axis on BM inflammation, we administered the CXCR4 receptor antagonist AMD3100 (5 mg/kg/day) intraperitoneally to 10-week-old control and MKO mice for 3 weeks. To determine whether AMD3100 can reduce infiltration of proinflammatory immune cells into the BM of MKO mice, we performed flow cytometry analysis to evaluate the proportions of diverse types of immune cells in the BM. AMD3100 induced a significant decrease in CD3+ T-cells of the BM in MKO mice, but it caused little change in the populations of BM CD4+ and CD8+ T-cells in either WT or MKO mice (Fig 6b and Supplementary Fig 7a). Moreover, the production of proinflammatory cytokines including IFN-γ, TNF-α, and IL-17A in CD4+ and CD8+ T-cells by BM cells was significantly reduced in the MKO mice treated with AMD3100 (Supplementary Fig 7b-e). Additionally, the expression of proinflammatory cytokines by the effector CD4+ and CD8+ T-cells was significantly decreased in the BM cells from MKO mice treated with AMD3100 (Fig 6c-h and Supplementary Fig 7f). These data indicate that CXCR4 antagonism attenuates BM infiltration by inflammatory T-cells in the MKO mouse model of muscular mitochondrial dysfunction.

**Figure 6.**
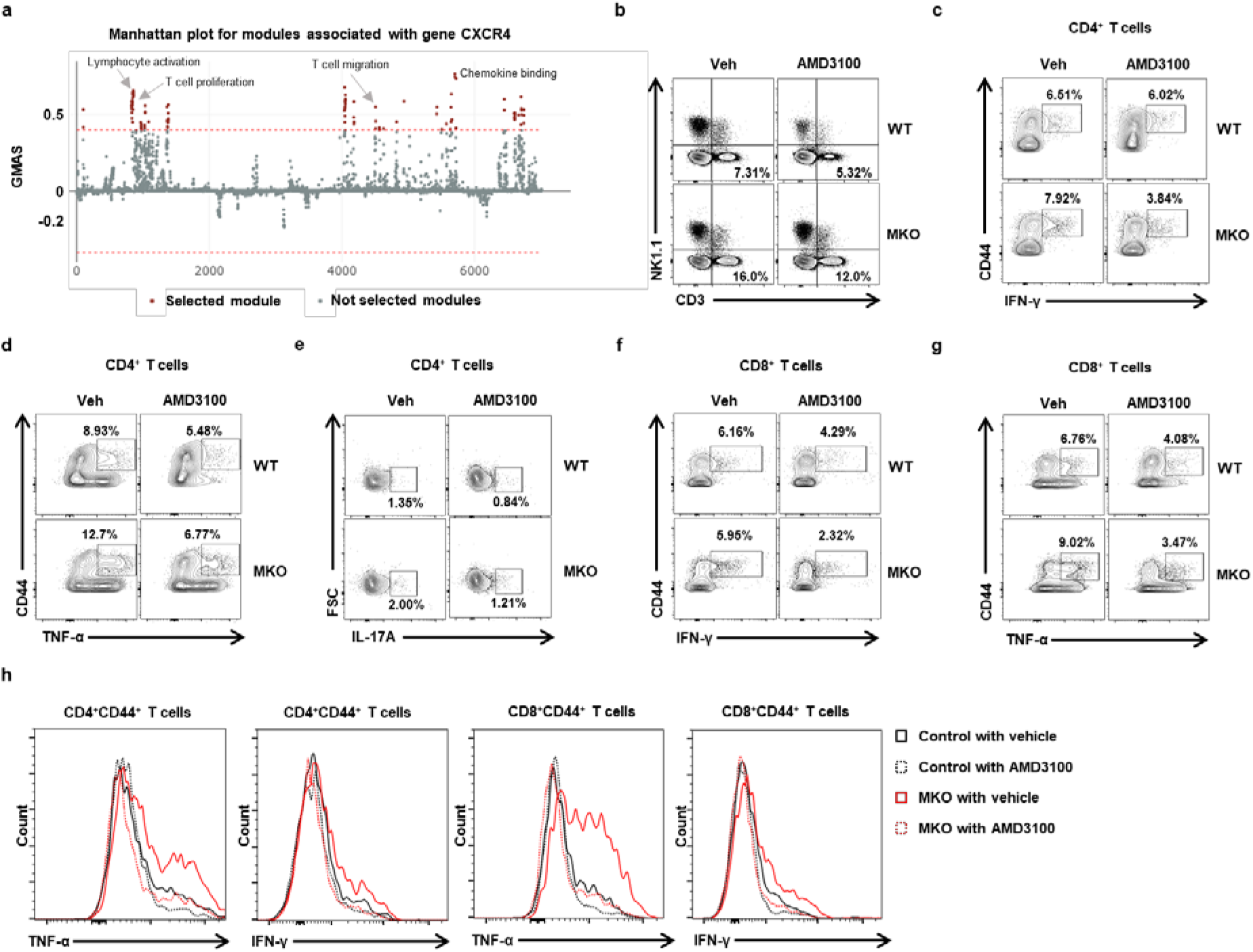
Treatment with CXCR4 antagonist reduces BM inflammation in MKO mice. (a) In the G-MAD analysis, CXCR4 associates with T-cell migration and proliferation, and chemokine-binding modules in mice. The threshold of significant gene-module association is indicated by the red dashed line. Modules are organized by module similarities. Known modules connected to CXCR4 are highlighted in red. GMAS, gene-module association score. (b) BM cells were subjected to flow cytometry for mature T-cells, natural killer, and natural killer T-cell phenotypes. (c,d) Populations of BM CD44+IFN-γ+ or CD44+ TNF-α+ among CD4+ T-cells from control and MKO mice treated with AMD3100 or vehicle at 14 weeks of age. (e) IL-17A production by BM CD4+ T-cells was analyzed by flow cytometry. (f,g) Populations of BM CD44+IFN-γ+ or CD44+ TNF-α+ among CD8+ T-cells from control and MKO mice treated with AMD3100 or vehicle at 14 weeks of age. (h) Production of IFN-γ or TNF-α by BM CD4+CD44+ or CD8+CD44+ T-cells of control and MKO mice treated with AMD3100 or vehicle at 14 weeks of age. Data are expressed as mean ± SEM. Statistical significance was analyzed by one-way ANOVA.

### Treatment with CXCR4 antagonists prevents bone loss in MKO mice

To investigate the effect of CXCR4 antagonism on bone loss in MKO mice, micro-CT analysis was conducted to assess BMD, BV, and cortical and trabecular bone architecture in the control and MKO mice treated with or without AMD3100. Consistent with previous results, MKO mice showed lower cortical and trabecular BV, BMD, trabecular number, and Tb.Th. compared with the controls (Fig 7a-c). AMD3100 significantly increased cortical and trabecular BV and trabecular number in MKO mice but not in the controls, without any changes in markers indicative of liver injury or perturbed lipid metabolism (Fig 7a-c and Supplementary Fig 8a-d). Taken together, these findings demonstrate that the CXCL12–CXCR4 axis is important for the BM inflammatory response and bone loss in the MKO mouse model with muscle dysfunction caused by reduced mitochondrial OxPhos.

**Figure 7.**
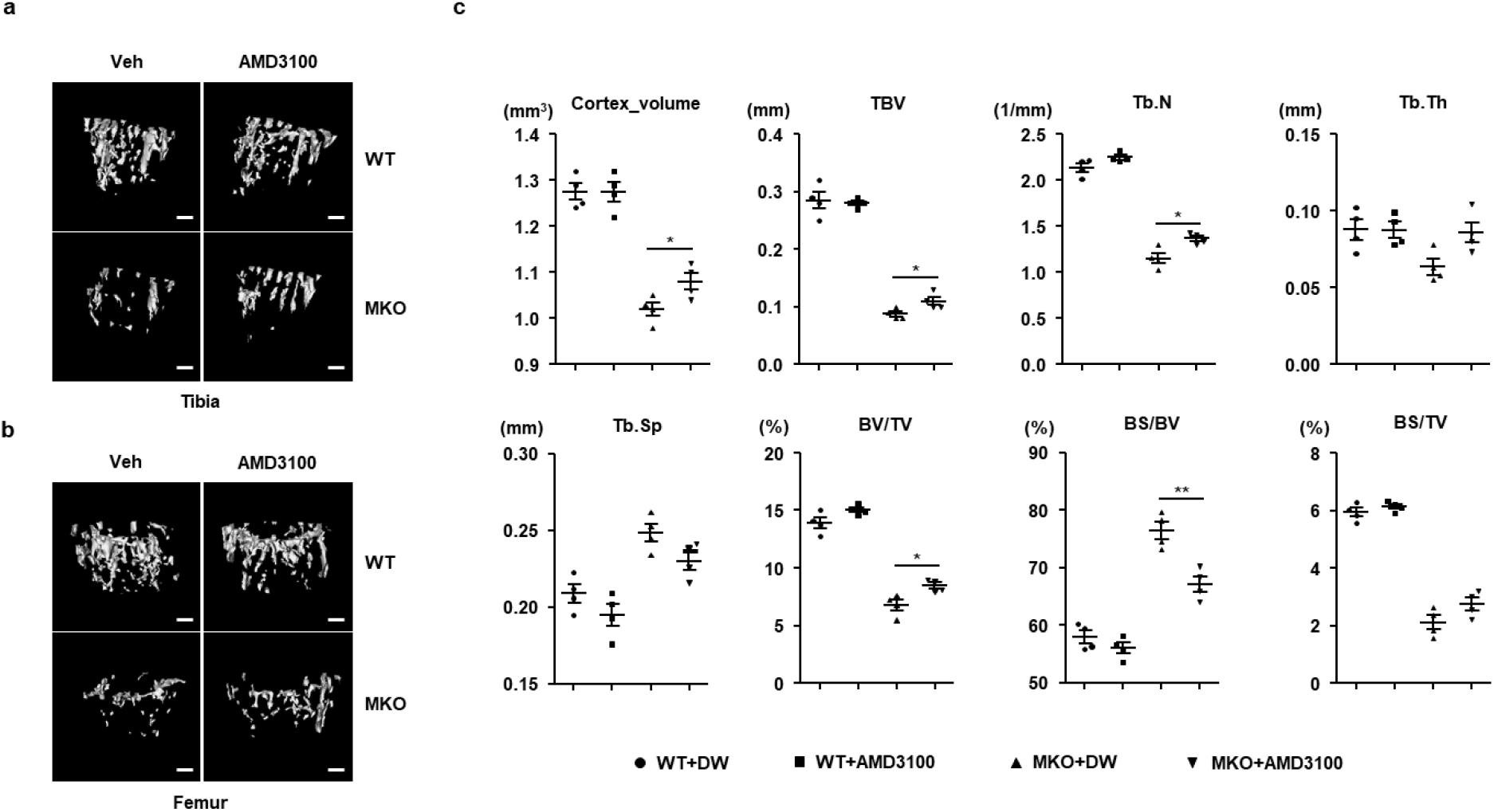
Treatment with AMD3100 attenuates bone loss in MKO mice. (a,b) At 9 weeks of age, control and MKO mice were injected intraperitoneally with AMD3100 (5 mg/kg, three times per week) for 3 weeks. Tibial and femoral trabeculae of control and MKO mice were measured by micro-CT. Scale bars, 250 μm. (c) Measurement of Tb.N., Tb.Th., BV/TV, BS/TV, BS/BV, Ct.V., Tb.Sp., and TBV in the tibiae from control and MKO mice using micro-CT analysis. Data are expressed as mean ± SEM. Statistical significance was analyzed by one-way ANOVA. *, *P* < 0.05 and **, *P* < 0.01 compared with the indicated group. DW, distilled water.

### Higher inflammatory response in the BM of hip fracture patients with lower body mass index

Muscle mitochondrial impairment is an important feature of pre-frailty development in humans ^21^. It is also well known that body mass index (BMI) has a positive correlation with muscle strength and mass in humans ^22^. As shown in Supplemental Figure 9A, grip strength was significantly lower in hip fracture patients with lower BMI compared with age- and gender-matched control subjects (Supplemental Table 1). We next investigated the immunophenotype of T-cells in the BM of hip fracture patients with lower or normal BMI (Fig 8a). To compare the different subsets of T-cells in the BM of patients with a lower BMI with those with a normal BMI, we evaluated the frequency of CD4+ and CD8+ BM T-cells expressing naïve/memory markers (CD45RA+/RO+). The patients with lower BMI had larger and smaller populations of CD8+ T-cells and CD4+ T-cells, respectively (Fig 8a and Supplementary Fig 9b). In the subset analysis of CD4+ and CD8+ T-cells, in both populations the proportion of CD45RA- CD45RO+ memory T-cells was significantly increased, while that of CD45RA+CD45RO-naïve T-cells was decreased in patients with lower BMI (Fig 8b-c, and Supplementary Fig 9c,d). Similarly to the role of T-cell senescence in chronic inflammatory conditions ^23^, the population of BM CD57+ senescent CD4+ and CD8+ T-cells was also larger in the patients with lower BMI (Fig 8d,e, and Supplementary Fig 9e). Furthermore, the expression of bone resorptive cytokines, including TNF-α and IL-17A, was markedly increased in the BM CD4+ and CD8+ T-cells of patients with lower BMI (Fig 8f-i). The expression of inflammation-related genes including *CXCL12*, *CD44*, *TNF*, and *IL17A* was also higher in the BM of patients with lower BMI (Supplementary Fig 9f-i). Moreover, serum levels of FGF21 and growth differentiation factor 15 (GDF15) trended higher in patients with lower BMI (Supplementary Fig 10a,b). Collectively, these data suggest that lower BMI and muscle function predict a larger population of proinflammatory and cytotoxic senescent T-cells in the BM of patients with hip fracture.

**Figure 8.**
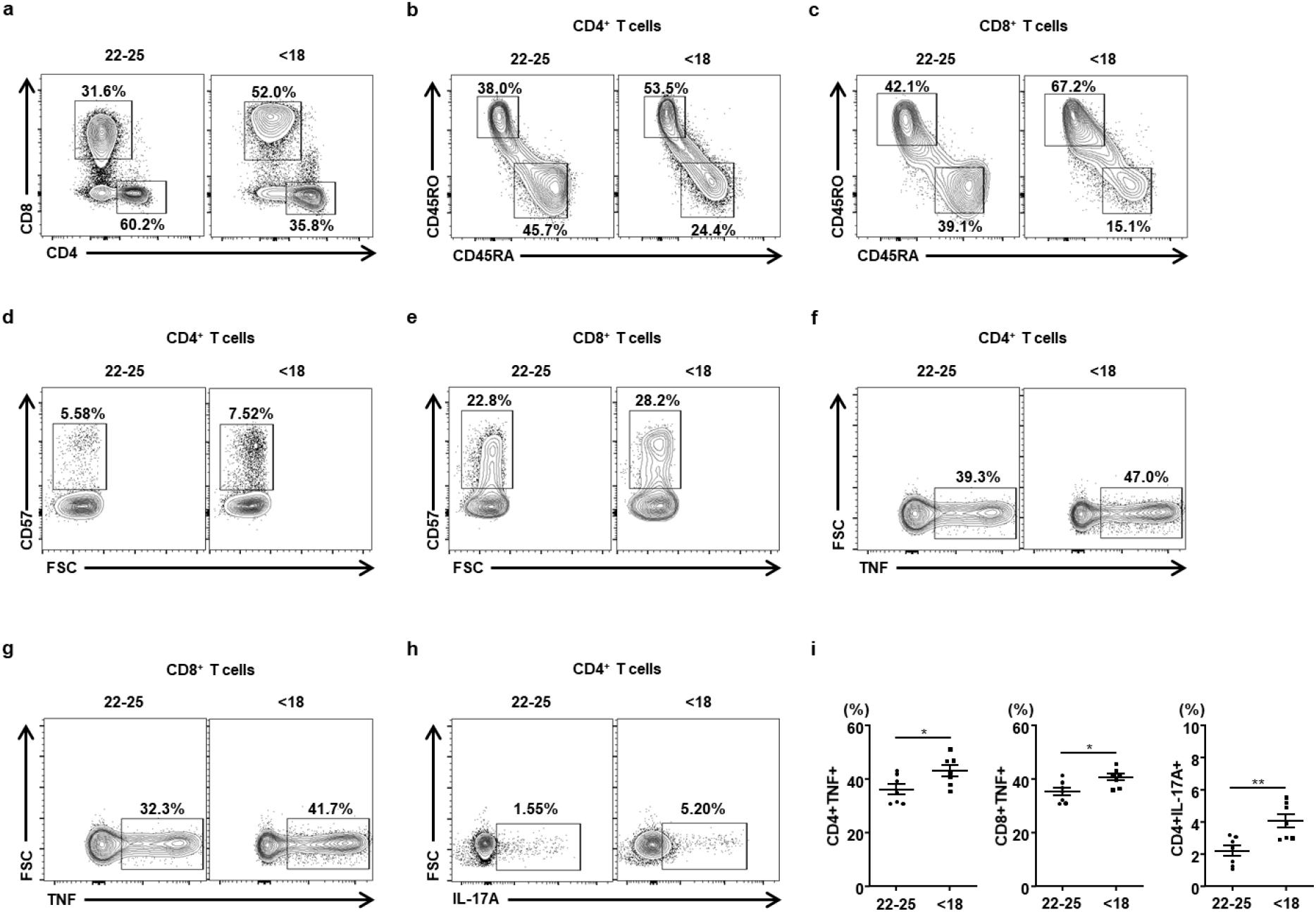
Immunophenotyping of BM T-cells in patients with hip fracture according to BMI. (a) Representative flow cytometry contour plots are presented for CD4 and CD8 expression by CD3+ T-cells in hip fracture patients with normal (22–25 kg/m^2^) and lower (<18 kg/m^2^) BMI. (b,c) Representative plots of CD45RO and CD45RA expression among CD4+ or CD8+ T-cells in the BM from hip fracture patients with a lower (<18 kg/m^2^) or normal (22–25 kg/m^2^) BMI (each *n* = 7). (d,e) CD57+ senescent population in CD4+ and CD8+ T-cells of the BM from hip fracture patients with a lower (<18) or normal (22–25) BMI (each *n* = 7). (f-h) Frequency of TNF-α- or IL-17A-producing cells among the BM CD4+ and CD8+ cells were evaluated by flow cytometry (each *n* = 7). (i) Statistical analysis of the population of TNF-α- or IL-17A-producing cells among the BM CD4+ and CD8+ cells in the two groups (*n* = 5/group). Data are expressed as mean ± SEM. Statistical significance was analyzed by unpaired t-tests. *, *P* < 0.05 and **, *P* < 0.01 compared with the indicated group.

## Discussion

In the present work, we investigated the impact of the mitochondrial OxPhos function in skeletal muscle on the maintenance of bone mass in mice. Our observations indicate that lower mitochondrial OxPhos function characterizes a reduction in muscle mass and physical activity, which contribute to the bone fragility of MKO mice. In addition, we report that mitochondrial stress in skeletal muscle induces BM inflammation, wherein the CXCL12–CXCR4 axis is involved in the migration and activation of T-cells. Antagonism of CXCR4 protects against the bone loss of the MKO mouse model with muscular mitochondrial dysfunction through attenuation of BM inflammation.

Mitochondria are important organelles regulating critical cellular processes in the physiology and pathology of skeletal muscle ^24^. Consistent with this, our mouse model with skeletal muscle-specific mitochondrial OxPhos dysfunction showed lower muscle mass and physical performance (Fig 1), making it suitable for use as a model of muscle atrophy. Moreover, it has been suggested that appendicular muscle mass and strength are closely associated with BMD ^25^. In line with this report, we have demonstrated that MKO mice also exhibit lower bone mass, as well as a reduction in cortical and trabecular BMD (Fig 2). Previous reports also indicate a positive correlation between mass and function of skeletal muscle and bone mass, which may be due to reduced physical loading or a diminished response to load. However, the biological mechanism of muscle dysfunction-induced bone loss and fragility remains to be determined. To this end, in the current study, we showed through an osteoimmunologic analysis of MKO mice that BM inflammation plays an essential role in muscle dysfunction-induced low bone mass and structural deterioration of bone tissue. Thus, it is possible that the improvement of muscle mitochondrial function by exercise training and/or diet may enhance bone mass and quality.

Inflammation is regarded as playing a causal role in bone loss in humans and mice with sex steroid deprivation. TNF-α-producing T-cells are increased in the BM of mice and humans through natural or surgical menopause, contributing to bone loss and fragility ^26–29^. In addition, estrogen deficiency induces high levels of serum IL-17 and promotes Th17 cell differentiation in ovariectomized mice ^30^. IL-17 is increased in patients with primary hyperparathyroidism and also mediates parathyroid hormone-induced bone loss ^31^. Moreover, systemic inflammation caused by sepsis produces osteoblast ablation and bone loss without any change in osteoclast function ^32^. In the current work, we show that, in MKO mice, muscular mitochondrial OxPhos dysfunction induces an increase in TNF-α- and IL-17A-producing T-cells of the BM. The hip fracture patients with a markedly lower BMI also exhibited a higher inflammatory response, as measured by proinflammatory cytokine production, in the BM. Collectively, the present findings indicate that BM inflammation is required for loss of muscle mass and function-mediated bone loss and fragility.

CXCR4 stimulates signal transduction for T-cell chemotaxis and gene expression via association with the T-cell receptor ^33^. CXCR4 is required for the migration of pathogenic CD4+ and CD8+ T-cells to the BM in a mouse model of aplastic anemia ^34^. In addition, CXCR4 overexpression by CD8+ T-cells increases their migration toward the BM microenvironment, leading to memory T-cell differentiation and production of effector cytokines including IFN-γ, TNF-α, and IL-2 ^35^. Adoptively transferred central memory T-cells accumulated more efficiently in BM cavities via their higher level of CXCR4 expression compared with naïve and effector T-cells ^36^. One of the most prominent features of the present work is the observation that CXCR4 antagonism protected MKO mice against BM inflammation and bone loss. Consistent with these findings, the CXCR4 antagonist AMD3100 has been shown to improve ovariectomy-induced bone loss by facilitating mobilization of hematopoietic progenitor cells ^37^. Moreover, treatment with vascular endothelial growth factor and AMD3100 can mobilize mesenchymal stem cells toward fracture healing, leading to bone formation in a delayed union osteotomy model ^38^. CXCL12, a chief ligand for CXCR4, is not only linked with disease severity of postmenopausal osteoporosis ^39^, but also acts as a pro-inflammatory factor in the progression of collagen-induced osteoarthritis by attracting inflammatory cells to joints and by activating osteoclasts ^40^. On the other hand, genetic disruption of CXCR4 enhances osteoclastogenesis and leads to increased osteolytic tumor growth in bone ^41^. Discrepancies in findings between different studies could be attributed to the use of different mouse or disease models and/or to the distinct role of CXCR4 in many different kinds of cell. Therefore, further investigation is required to establish the role of CXCR4 in the diverse context of bone loss using a cell type-specific *Cxcr4* knockout mouse model.

In this study, we analyzed two kinds of RNA sequencing data derived from EDL muscle and whole BM cells from control and MKO mice. We observed that mitochondrial stress via OxPhos dysfunction increases local expression and serum levels of FGF21, which do not have any effects on bone mass and quality in MKO mice (Fig 3). On the other hand, GSEA indicated that BM cells from MKO mice showed prominent expression differences in inflammatory response and adipogenesis gene sets compared with the controls (Fig 5). Ectopic fat, defined as storage of triglyceride in tissues other than adipose tissue, is associated with metabolic deterioration in humans as well as rodents. Although the pathogenesis of ectopic fat deposition is largely unknown, free fatty acids released by adipocyte hypertrophy and inflammatory response are important in the development of ectopic fat-induced organ dysfunction in liver, skeletal muscle, and heart ^42^. Excessive fat deposition in non-adipose tissues recruits immune cells including macrophages and activated T-cells, thereby promoting chronic inflammation in the metabolic organs ^43^. Several investigations have revealed that BM is also susceptible to fat deposition through old age, menopause, obesity, anorexia nervosa, and weight loss surgery ^44–47^. Although multiple studies have investigated ectopic adipocyte accumulation in BM cavities, our understanding of its role in the inflammatory response in BM is incomplete. Ectopic fat accumulation in BM is associated with activation of immune cells and bone loss ^48^. Taken together, these data suggest that muscular mitochondrial dysfunction-mediated bone loss is caused by the change in the BM microenvironment rather than the change in mitokine secretion.

In conclusion, mitochondrial OxPhos function in skeletal muscles play a pivotal role in the regulation of BM inflammation and bone loss in mice. Inhibition of BM inflammation by a CXCR4 antagonist attenuates bone loss in a mouse model with muscle mitochondrial dysfunction. However, the human relevance of muscular mitochondrial OxPhos dysfunction and BM inflammation in the regulation of skeletal homeostasis needs to be clarified.

## Methods

### Animals

To generate skeletal muscle-specific *Crif1* deficiency in mice, floxed *Crif1* mice were crossed with *Mlc1f*-*Cre* mice on a C57BL/6 background (provided by S.J. Burden, New York University, Baltimore, MD, USA). MKO mice were crossed with a global *Fgf21* knockout on a C57BL/6 background (a kind gift from N. Itoh, Kyoto University Graduate School of Pharmaceutical Sciences, Kyoto, Japan) to generate MKFO mice as a mitokine double knockout mouse model. All animal experiments used male mice that were maintained in a controlled environment (12 h light/12 h dark cycle; humidity, 50–60%; ambient temperature, 22 ± 2°C) and fed a normal chow diet. All experimental procedures involving mice were conducted in accordance with the guidelines of the Institutional Animal Care and Use Committee of Chungnam National University School of Medicine (CNUH-017-A0048, Daejeon, Republic of Korea).

### Human subjects

Patients with hip fracture were recruited from the Chungnam National University Hospital between October 2019 and February 2020. The participants were divided into two age- and gender-matched groups as follows: BMI, 22–25 (n = 7) and BMI <18 (n = 7). Patients with any of the following conditions were excluded from the study: rheumatoid arthritis, neuromuscular disorder, chronic kidney disease, and mineral and bone disorder. Patients with diseases that affect bone metabolism (or those taking drugs that affect bone metabolism), history of any malignant or inflammatory disease, and past hormone replacement therapy were also excluded. Hand grip strength was measured using an electronic hand dynamometer (Lavisen, Namyangju, South Korea). Grip strength of the dominant hand was measured only once in a sitting posture with 0° shoulder angle, 90° elbow angle and neutral wrist angle. Lymphocytes were isolated from the BM cells of the enrolled patients and stored at −180°C in liquid nitrogen prior to flow cytometry analysis. Whole BM cells were used for real-time PCR analysis. This human study was reviewed and approved by the Institutional Review Board of Chungnam National University Hospital (CNUH 2019-10-065), according to the standards of the Declaration of Helsinki. Written and oral informed consent, documented by the Department of Internal Medicine of Chungnam National University Hospital in South Korea, was obtained from all of the participants prior to their inclusion in the study.

### Flow cytometry analysis

Isolated BM cells were filtered through a 70 μm cell strainer, washed in phosphate-buffered saline (PBS), and resuspended in a 40% Percoll (GE Healthcare, Chalfont St Giles, UK) gradient. The cell suspension was centrifuged at 2,400 rpm for 30 min at 4°C. Then, the cells were incubated with directly fluorochrome-conjugated monoclonal antibodies for 40 min at 4°C. The antibodies used in this study are listed in Supplemental Table 2. For blocking non-specific antibody binding, cells were pre-incubated with anti-mouse CD16/32 mouse Fc blocker (BD Biosciences, San Jose, CA, USA) prior to staining with the antibodies. For intracellular staining, BM cells were stimulated with Cell Stimulation Cocktail including phorbol myristate acetate, ionomycin, brefeldin A, and monensin (eBioscience, San Diego, CA, USA) for 5 h. The cells were fixed and permeabilized using a Fixation/Permeabilization Buffer kit (eBioscience, San Diego, CA, USA), and then washed and resuspended in 1% formaldehyde, and further stained for intracellular cytokines with anti-IFN-γ-PE-Cy7, anti-TNF-α-APC, or anti-IL-17A-APC. Multicolor flow cytometry was performed using a BD LSRFortessa flow cytometer (BD Biosciences), and the data were analyzed using FlowJo software (Tree Star, Ashland, OR, USA). Results are expressed as cell frequency (%).

### Immunoblots

Tissues and cells were lysed using 2% sodium dodecyl sulfate (SDS) with 2 M urea, 10% glycerol, 10 mM Tris-HCl (pH 6.8), 10 mM dithiothreitol, and 1 mM phenylmethylsulfonyl fluoride. The lysates were centrifuged, and the supernatants were separated by SDS-polyacrylamide gel electrophoresis (PAGE) and blotted onto a nitrocellulose (NC) membrane. After blocking with 5% skimmed milk, the membrane was analyzed using specific antibodies and visualized by enhanced chemiluminescence using WesternBright ECL Spray (Advansta, Menlo Park, CA, USA). Proteins were detected by immunoblotting with antibodies listed in Supplemental Table 2. A horseradish peroxidase-conjugated goat anti-rabbit IgG (Enzo Life Sciences, Farmingdale, NY, USA) secondary antibody was used for visualization. Signals were obtained using an Odyssey imager and Image Studio Software (LI-COR Biosciences, Lincoln, NE, USA). Serum levels of FGF21 were measured using an ELISA kit (R&D Systems, Minneapolis, MN, USA).

### BN-PAGE

To isolate mitochondria from the EDL and gastrocnemius of 14-week-old control and MKO mice, samples were homogenized in isolation buffer (210 mM mannitol, 70 mM sucrose, 1 mM EGTA, and 5 mM HEPES, pH 7.2) using a Teflon-glass homogenizer. The homogenized tissues were centrifuged at 600 × g for 5 min at 4°C, and the supernatant was re-centrifuged at 17,000 × g for 10 min at 4°C. The isolated mitochondrial fraction was supplemented with 0.5% (w/v) n-dodecyl-β-D-maltoside and assessed for OxPhos complex content using a Native PAGE Novex Bis-Tris Gel system (Invitrogen, Carlsbad, CA, USA). The separated proteins were transferred to polyvinylidene fluoride membranes, which were incubated overnight at 4°C with an anti-OxPhos antibody cocktail (Invitrogen, #45-8099, #45-7999), and were analyzed using the Western Breeze Chromogenic Western Blot Immunodetection Kit (Invitrogen).

### SDH staining

Transverse 12 μm muscle sections were mounted on Super Frost microscope slides (Thermo Fisher Scientific, Waltham, MA, USA). Muscle fiber type-specific diameter measurements were obtained using 12 μm-thick SDH-stained cross-sections at 14 weeks of age. Sections were outlined with a PAP pen (Research Products International) and incubated in buffer solution (20 mM phosphate buffer, 7.5% sucrose, 0.027% sodium succinate, and 10 mg nitrobluetetrazoleum) for 45 min at room temperature. The sections were dehydrated, and then briefly rinsed in 30%, 60%, and 90% acetone in distilled water in ascending and descending order, rinsed in distilled water, air-dried, and cover-slipped using VectaMount (Vector Labs).

### Micro-CT analysis

Micro-CT was performed on vertebrae and long bones using an eXplore Locus SP scanner (GE Healthcare, London, Canada) with 8 μm resolution. All bone morphometric parameters were calculated three‐dimensionally with eXplore MicroView version 2.2 (GE Healthcare), which was used for measuring BV, total volume (TV), percent BV (BV/TV), bone surface (BS), bone surface density (BS/TV), Tb.Th., and Tb.Sp. Bone parameters and density were analyzed at the region between 0.7 and 2.3 mm below the growth plate of the distal femur. Cancellous bone was analyzed in the distal area extending proximally 1.75 mm from the end of the primary spongiosa. The half the height of the bone with half the width of the 5th lumbar vertebral body was used to analyze cancellous spine. All bone micro-CT nomenclature follows the guidelines of the American Society for Bone and Mineral Research (ASBMR).

### Bone histological and morphological analysis

Mice were euthanized at 14 weeks and the bones were removed and fixed in 4% paraformaldehyde (Biosesang, Seongnam, korea) at 4°C overnight. To evaluate dynamic histomorphometry, mice were injected intraperitoneally with alizarin red (Sigma-Aldrich, Dorset, UK, 30 mg/kg) and calcein (Sigma-Aldrich, Dorset, UK, 10 mg/kg) 8 and 3 days prior to CO_2_ asphyxiation, respectively. For bone histological analysis, femurs and tibias of mice were harvested, skinned, and fixed in 4% paraformaldehyde overnight at room temperature. The samples were then dehydrated in ethanol solution and decalcified in 10% ethylenediaminetetraacetic acid (Sigma-Aldrich, Dorset, UK) for 2 weeks at room temperature. The buffer was changed every 3–4 days until complete decalcification. The tissues were embedded in paraffin, and 4 μm sagittal-oriented sections were prepared and stained with H&E and TRAP for histological analysis using standard protocols. For von Kossa staining, undecalcified bones were embedded in methyl‐methacrylate (Sigma) and sectioned at a thickness of 6 μm as previously described ^49^. Bone histomorphometric analysis was performed with the Bioquant Osteo II program (Bio‐Quant, Inc., Nashville, TN, USA). All bone histomorphometry nomenclature follows the guidelines of the ASBMR ^50^.

### Mouse grip strength and wire hanging test

Experiments were performed using a digital force-gauging apparatus (GS 5000; Borj Sanat, Iran). Mice were allowed to grasp the pull bar with fore limbs, and then they were gently pulled parallel away from the bar by the tail until the forelimbs released the bar. Mice were not trained before testing. The maximum force prior to release of the mouse’s paw from the bar was recorded. The test was repeated five times, and the average value of five consecutive measurements was reported as the mouse’s grip strength. The wire hanging test was conducted to assess motor function and neuromuscular grip strength. The mouse was placed on a cross-grip wire rack, which was then turned upside down 20 cm above a cage filled with soft bedding, after which hanging time was recorded. The average latency to fall of four trials was calculated for each animal. The maximum hanging time was used in the analyses.

### RNA sequencing

Total RNA was prepared from EDL and BM cells obtained from 10-week-old control and MKO mice (*n* = 2 for each) using TRIzol reagent. The integrity of the total RNA was assessed using an Agilent 2100 Bioanalyzer System (Agilent Technologies, Loveland, CO, USA) and an Agilent RNA 6000 Nano Kit (Agilent Technologies, Loveland, CO, USA). The library was prepared using a TruSeq 3000/4000 SBS Kit, v3. It preprocessed the raw reads from the sequencer to remove results from low-quality RNA or artifacts such as adaptor sequences, contaminant DNA, and PCR duplicates. The quality of the data produced is determined by the phred quality score of each base. The FastQC quality control tool gives a box plot of average base quality per cycle, and a phred quality score of 20 means that the assignment to that base is 99% accurate. Generally a phred score ≥20 is good quality, and those in the present study were ≥30 was 97.19%. The obtained reads were mapped to a reference Mus musculus (mm10) genome using HISAT2 v2.0.5. HISAT uses two types of index for alignment (a global, whole-genome index and tens of thousands of small local indexes). These aligned reads were then assembled from known genes/transcripts using a reference gene model in StringTie v.1.3.3b. Transcript frequencies were quantified as normalized values, taking into account transcript length and depth of coverage. Relative transcript abundance was expressed as fragments per kilobase of transcript per million fragments mapped (FPKM), and FPKM values ≤0 were excluded. One was added to each FPKM value for filtered genes, the filtered data were log2-transformed, and quantile normalization was applied. Differentially expressed gene (DEG) analysis was performed using FPKM value. Genes with a fold-change >2 and an independent t-test *P*-value <0.05 were extracted from the results of the DEG analysis. A heatmap was produced by color-coding standardized log gene expression levels (mean, zero; standard deviation, one) using R 3.5.1 available at http://www.r-project.org. Probe sets are shown as hierarchically clustered by similarity, based on Euclidean distance and the Ward aggregation algorithm.

### RNA sequencing analysis using bioinformatics tools

DEGs were then subjected to hierarchical clustering and phenotype ontology using Network2Canvas (http://maayanlab.net/N2C/). Phenotype categories were visualized on the grid according to gene-list similarity, with enriched categories being indicated by circles. GSEA (http://www.broadinstitute.org/gsea) was performed on transcriptome data from the BM cells from control and MKO mice. Bioinformatic analysis was carried out with R package v3.2.5, available at http://www.r-project.org. A heatmap was produced by color-coding standardized log gene expression levels (mean, zero; standard deviation, one). Probe sets are shown as hierarchically clustered by similarity, based on Euclidean distance and the Ward aggregation algorithm. We also used G-MAD in GeneBridge tools (available at http://systems-genetics.org, an open resource), which uses expression data from large-scale cohorts to propose potential functions of genes and allows the annotation of gene function ^17^.

### Treatment with AMD3100 in vivo

Control and MKO mice (9 weeks of age) were injected intraperitoneally with 5 mg/kg PBS or AMD3100 (Sigma-Aldrich #A5602; St. Louis, MO, USA; Sigma Aldrich.com) three times a week for 3 weeks. At the end of treatment, the mice were sacrificed, and blood samples were collected for measuring proinflammatory cytokines and bone turnover markers. Femurs and tibiae were removed, fixed with 4% paraformaldehyde in PBS solution (pH 7.4) for 16 hr, and then stored at 4°C in 80% ethanol prior to measurement of BMD using micro-CT.

### Statistical analysis

Results are expressed as mean values ± standard error of the mean (SEM). Data were analyzed using Prism (version 8, GraphPad Software Inc., San Diego, CA, USA). A 2-tailed, unpaired t-test with Welch’s correction was used to assess statistical significance between two groups. A one-way analysis of variance (ANOVA) with Bonferroni’s correction for multiple comparisons was used to examine differences between more than two groups. A *P*-value <0.05 was considered statistically significant.

## Author contributions

MS and HSY designed the studies. JWT, JSM, HTN, HYL, JTK, JYC, XC, and HSY performed the research and analyzed the data. HKC, DR, JYC, MT, and TS analyzed the data. HSY and JWT wrote the manuscript.

## Acknowledgments

This work was supported by the Basic Science Research Program, through the National Research Foundation of Korea (NRF), funded by the Ministry of Science, ICT, and Future Planning, Korea (NRF-2018R1C1B6004439, NRF-2019M3E5D1A02068575) and the CNUH Research Fund, 2018. MS and SKK were also supported by the NRF (NRF-2017K1A1A2013124 and NRF-2017R1E1A1A01075126).

---

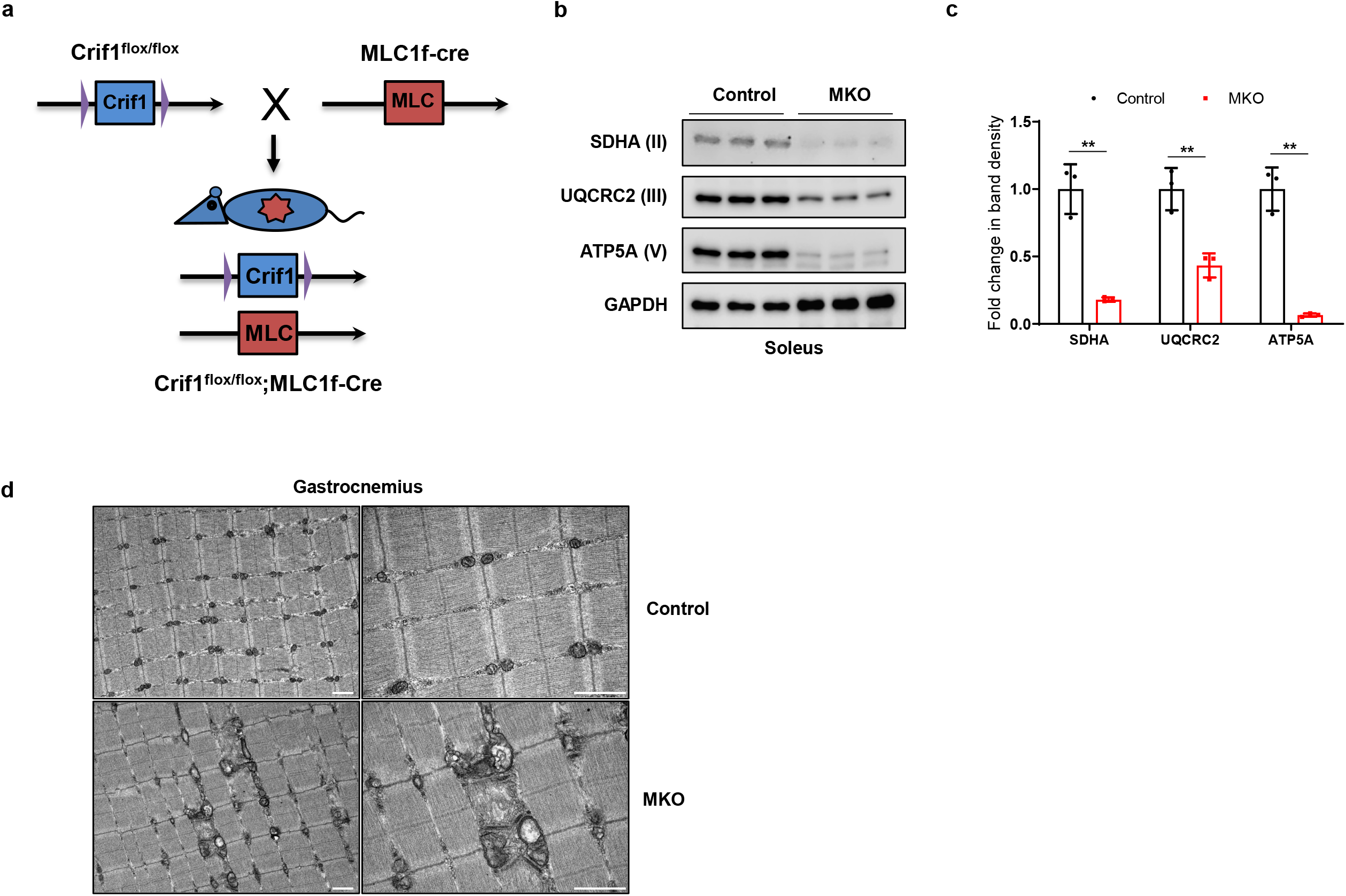

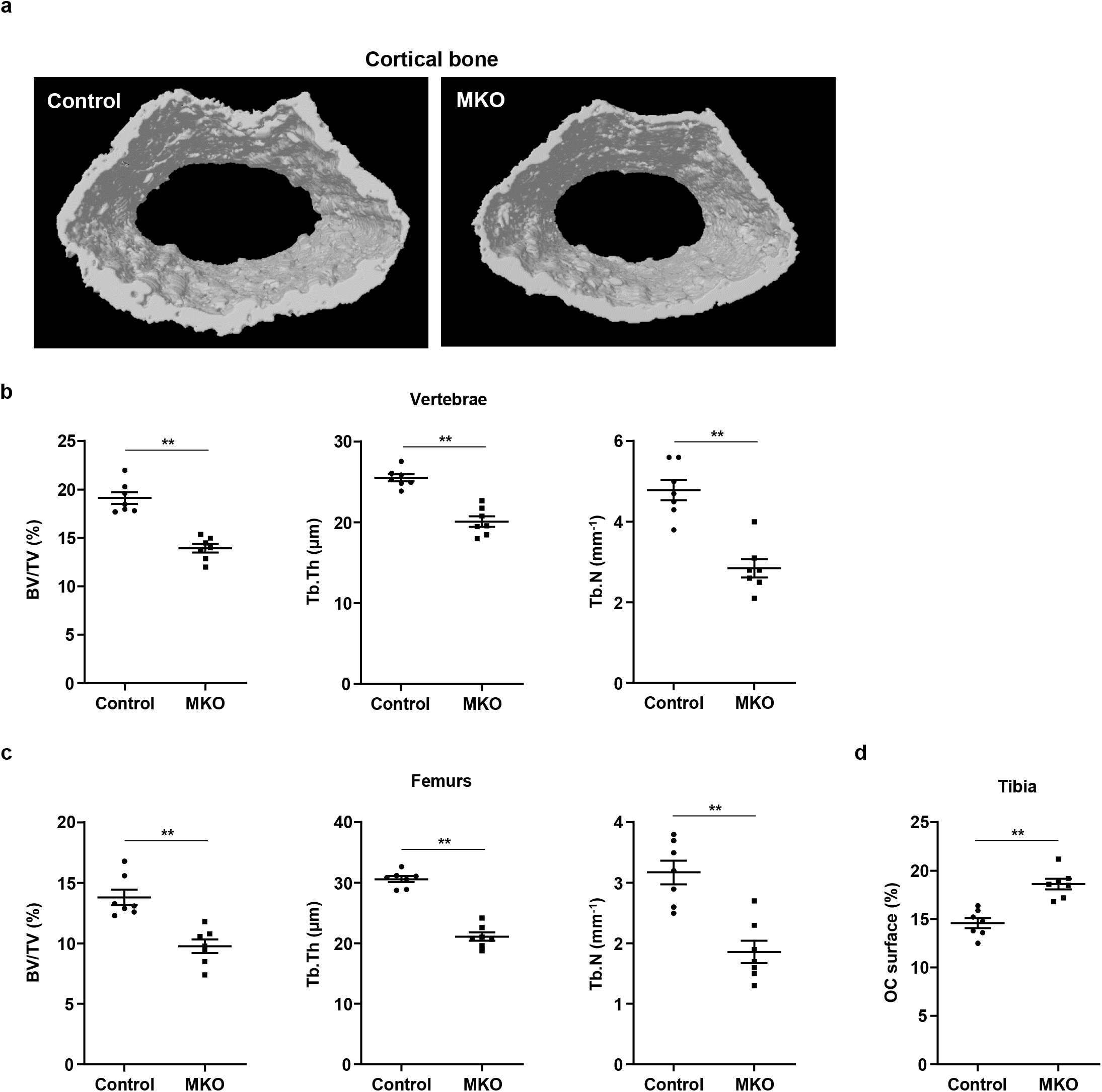

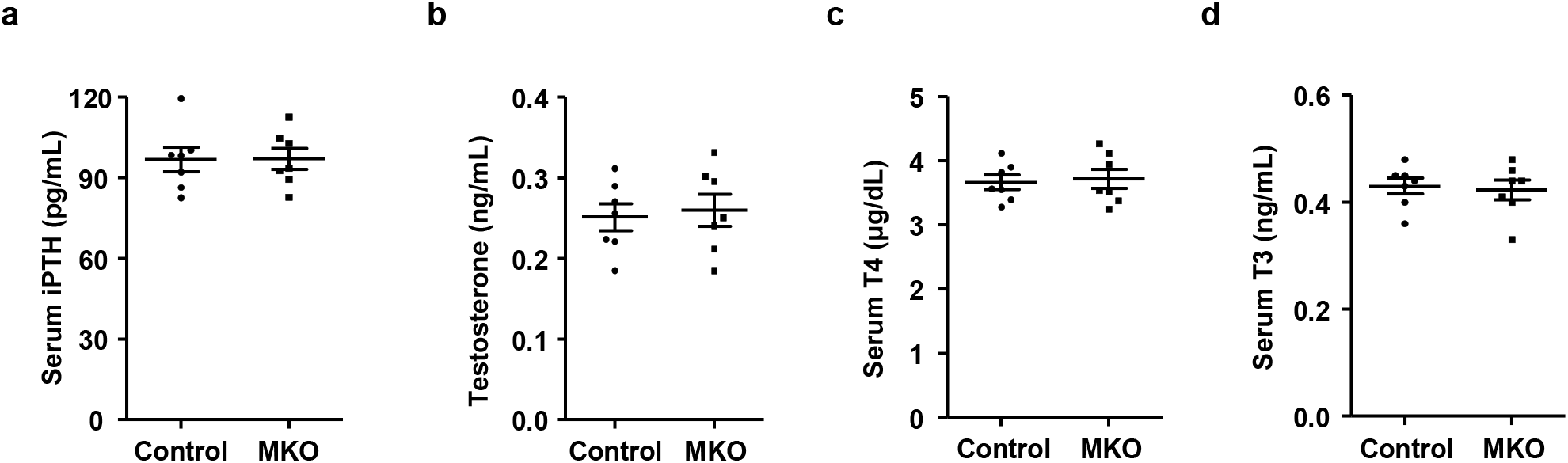

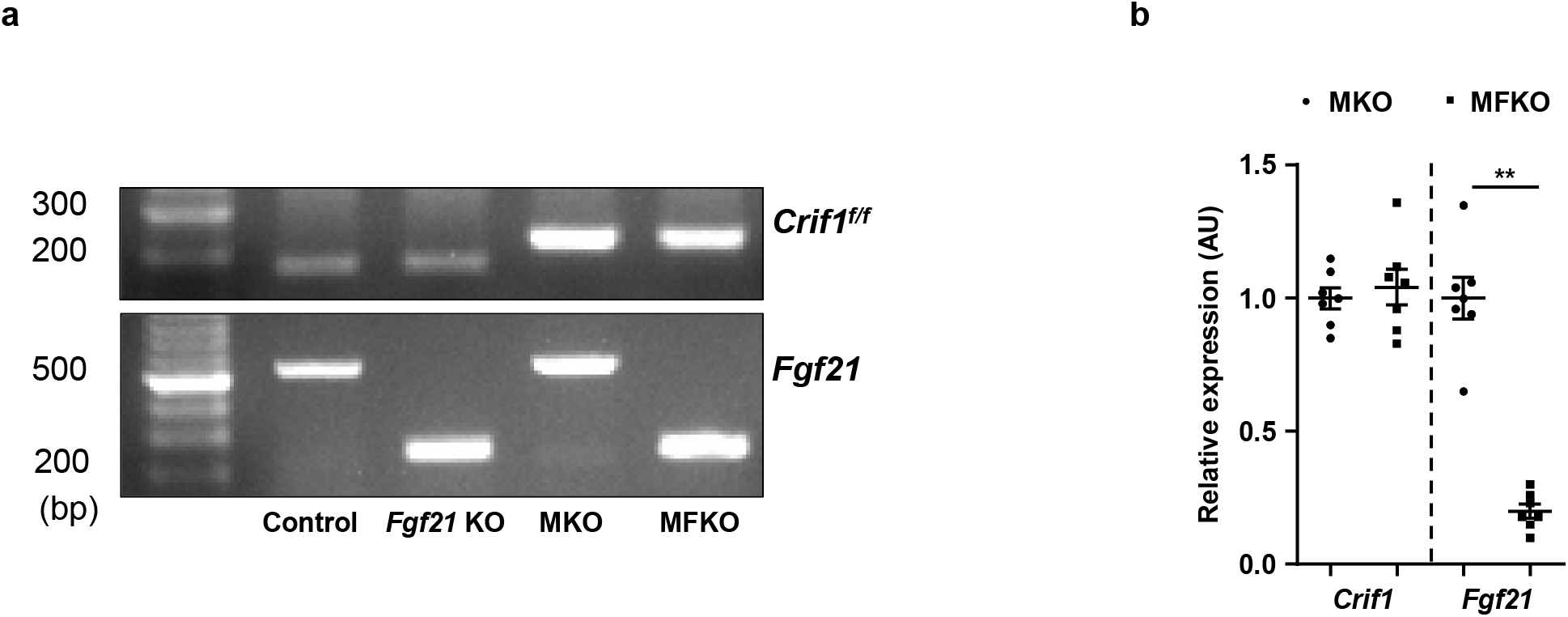

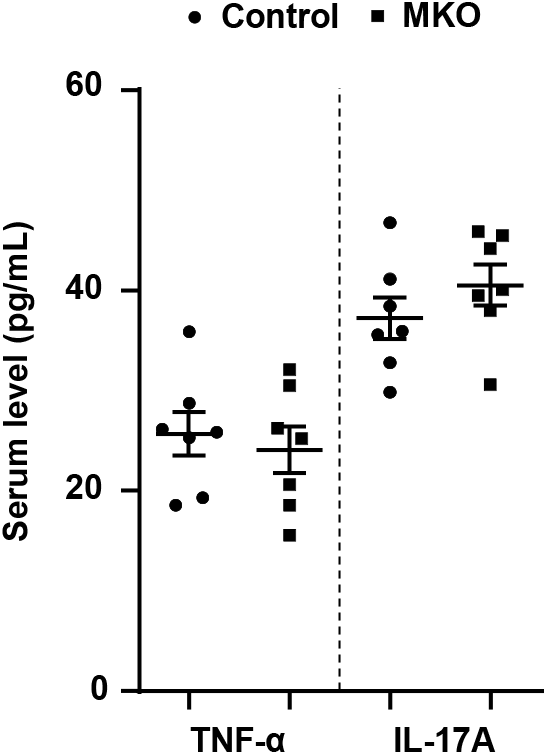

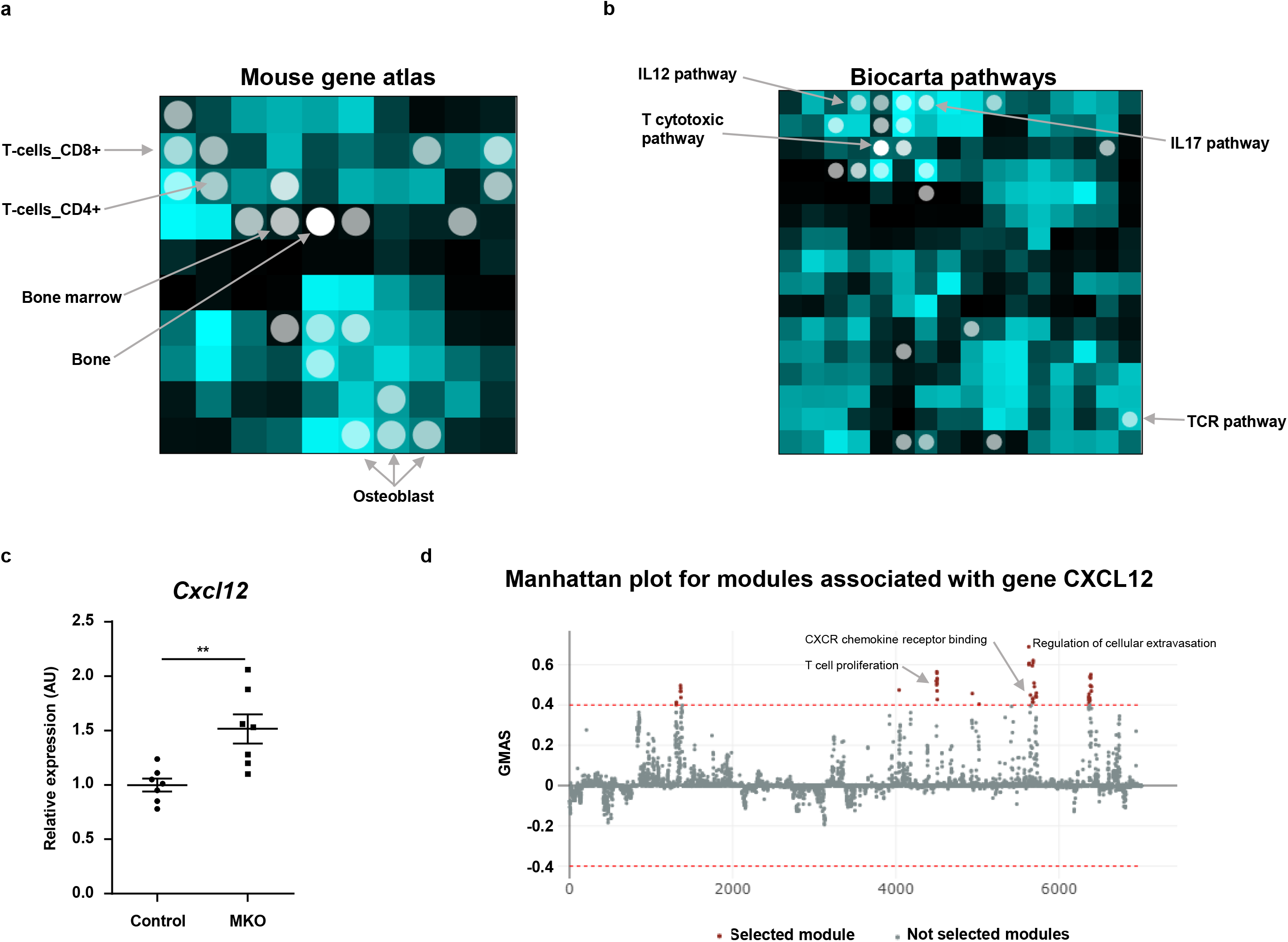

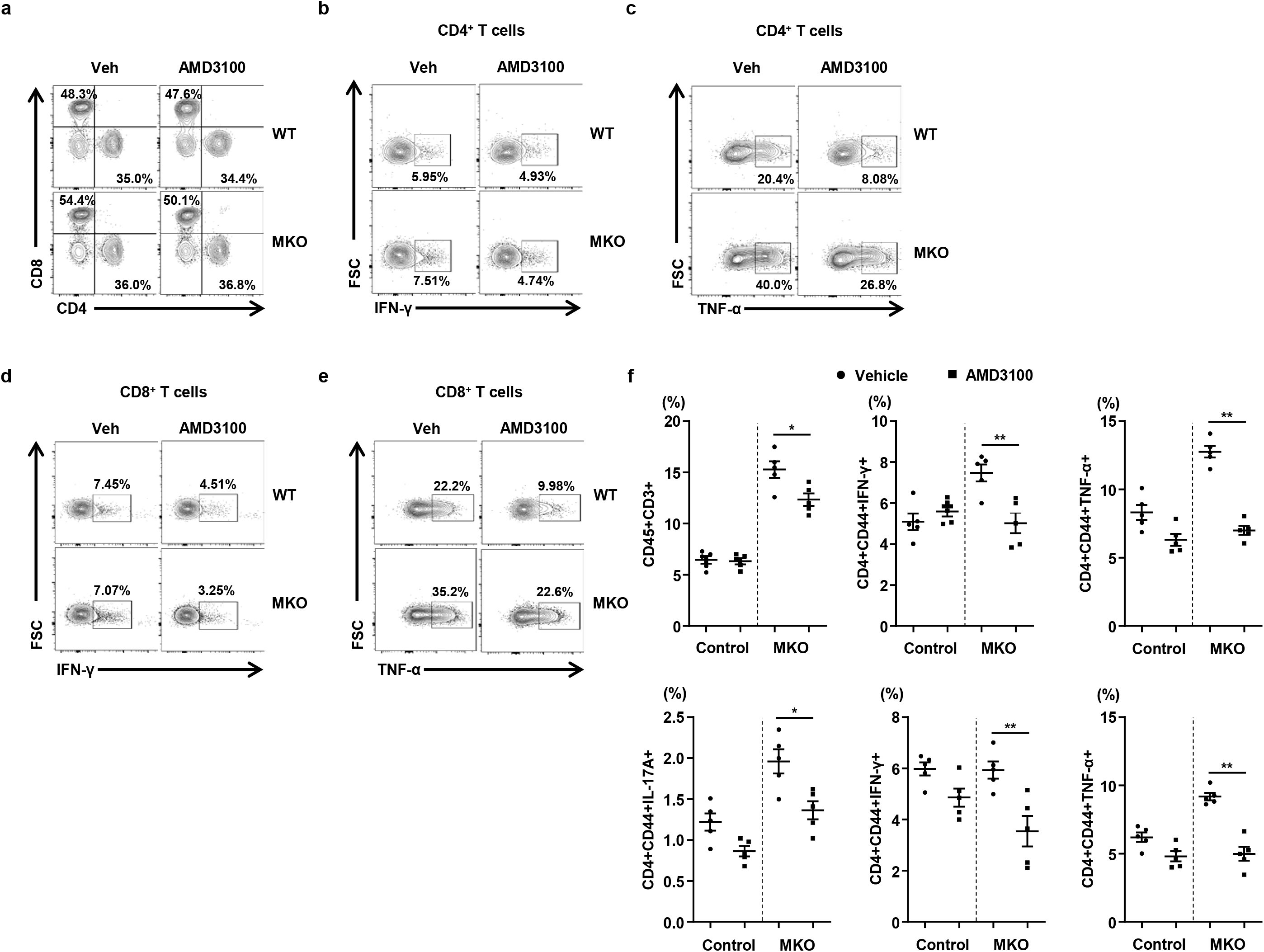

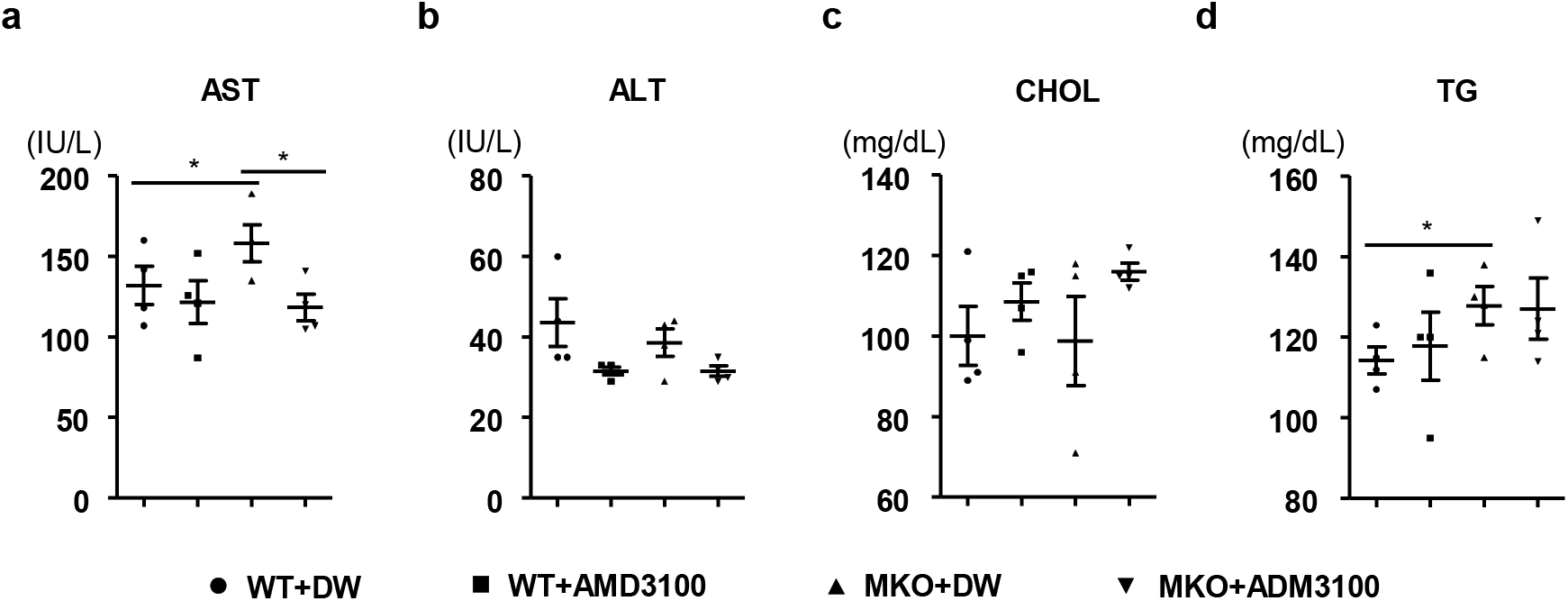

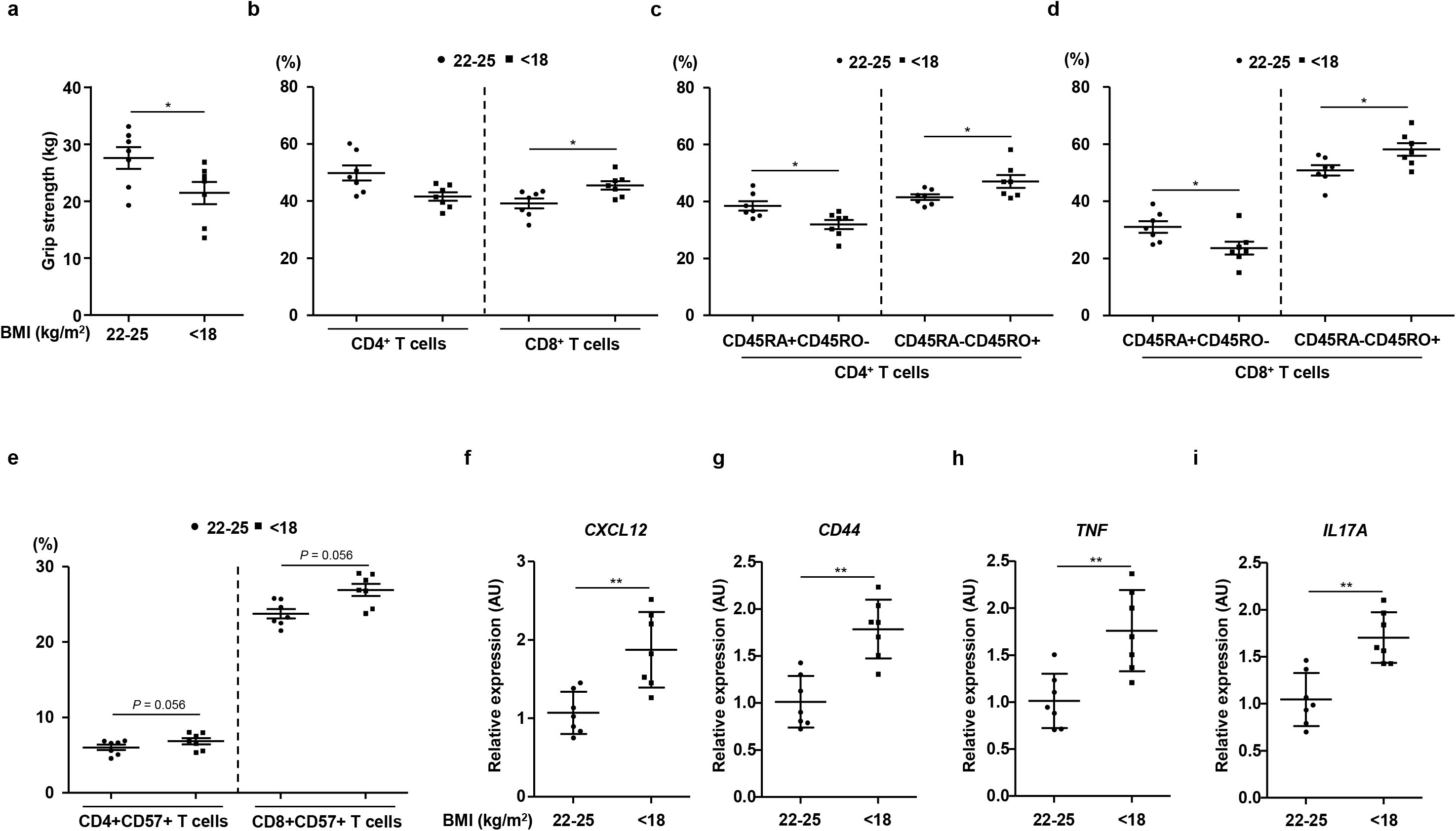

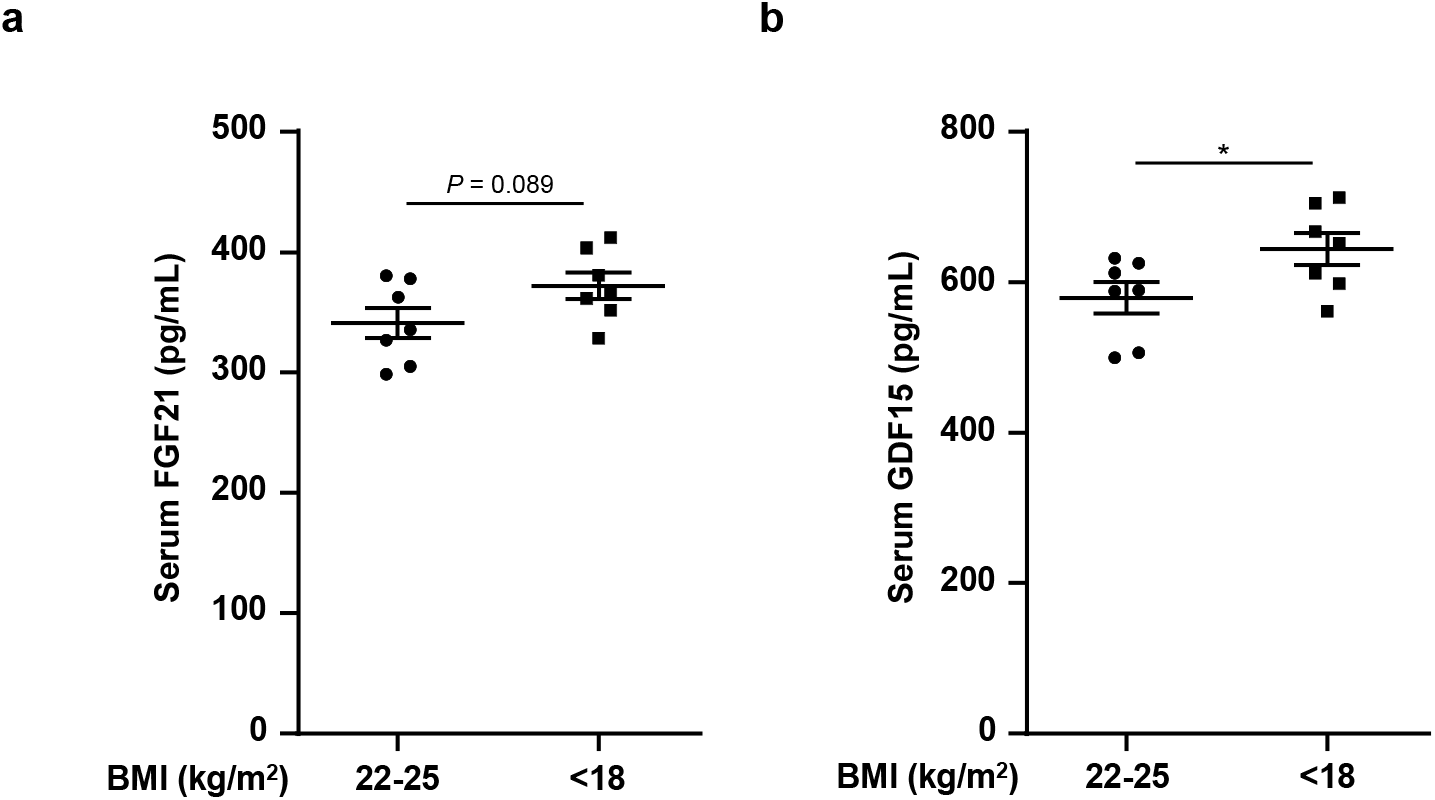

