## Supporting information for "Skeletal muscle mitochondrial dysfunction in mice is linked to bone loss via the bone marrow immune microenvironment"

***Conflict of Interest Statement***: The authors have declared that no conflict of interest exists.

***Key words***: mitochondria, inflammation, bone marrow, bone loss

***Word count***: 6,091 (Introduction, Results, Discussion, and Methods)

***Number of figures and tables***: 8

*Corresponding author: Hyon-Seung Yi, Research Center for Endocrine and Metabolic Diseases, Chungnam National University School of Medicine, Daejeon 35015, Korea. **Phone**: +82-42-280-6994; **Fax**: +82-42-280-6990; **E-mail**:

**Supplemental methods**

*Serum measurements.* Blood samples were collected by cardiac puncture of mice under general anesthesia. Samples were centrifuged at 10,000 rpm for 5 min, and the supernatant was used for assaying triglycerides, total cholesterol, alanine aminotransferase, and aspartate transaminase using a DRI-CHEM 4000i automated system (Fujifilm, Tokyo, Japan). Serum levels of TNF-α and IL-17A were determined using a specific enzyme-linked immunosorbent assay (ELISA; TNF-α, BD Bioscience, NJ, USA; IL-17A, Sigma-Aldrich, St. Louis, MO, USA) following the manufacturer's protocol. C-telopeptide (CTX), procollagen type 1 N propeptide (P1NP), testosterone, T3, and T4 were measured by ELISAs. Serum levels of T3, T4, and testosterone were measured using an ELISA kit (T3 and T4, Merck-Millipore, Darmstadt, Germany; testosterone, ALPCO, Salem, NH, USA). Serum parathyroid hormone (PTH) levels were measured using the mouse intact PTH ELISA kit (Immutopics International, San Clemente, CA, USA). In human subjects, serum FGF21 and GDF15 levels were assayed with a commercial ELISA kit (R&D Systems, Minneapolis, MN, USA) in accordance with the manufacturer's protocol.

*Preparation of bone marrow cells from mice and humans.* Control and MKO mice (14 weeks of age) were anesthetized by intraperitoneal injection of sterile avertin (tribromoethanol: 200 mg/10 ml/kg), and then tibias and fibulas were removed. The bone marrow (BM) cells were flushed from the medullary cavities of the excised bones and suspended in RPMI-1640 medium containing 10% fetal bovine serum

(FBS). Human BM cells were isolated from patients who underwent hip arthroplasty at the Chungnam National University Hospital between October 2019 and February 2020. The BM cells were extracted from the femoral neck cutting area without bacterial contamination, and were subjected to flow cytometry and real-time PCR analysis. The demographic and clinical characteristics of the patients with a body mass index (BMI) of $<18$ kg/m$^2$ and 22–25 kg/m$^2$ that were included in this study are shown in Supplemental Table 1.

*RNA extraction and real-time PCR analysis.* Total RNA was extracted from the BM cells using TRIzol reagent (Life Technologies, Eugene, OR, USA). Complementary DNA (cDNA) was synthesized from total RNA using M-MLV reverse transcriptase and oligo-dT primers (Invitrogen, Carlsbad, CA, USA). Specific sequences were amplified from each cDNA sample using SYBR Green PCR Master Mix (Applied Biosystems, Foster City, CA, USA) and specific primers (Supplemental Table 3 and 4) using a 7500 Real-Time PCR System and Software, v2.0.6 (Applied Biosystems). The comparative Ct method was used to determine relative expression, with 18s ribosomal RNA as the reference gene.

*Measurement of bone mineral density in human subjects.* Bone mineral density (BMD) was measured in the lumbar spine (1st–4th lumbar vertebrae) and femoral neck of the participants with hip fracture using dual energy X-ray absorptiometry with a Discovery (Hologic Inc., Marlborough, MA, USA) scanner. All BMD scans were conducted by well-trained examiners using standardized procedures following the manufacturer's recommended protocols. The lumbar BMD of 30 patients was measured with two consecutive measurements per patient. Any scans comprising metal or other attenuating material in the region of interest, as well as any scans of poor quality were discarded. The precision error of the lumbar BMD measurement was 1.4%, which was lower than the minimum acceptable precision error of 1.9% for the lumbar spine.

*Rotarod test of coordination*. Mice were trained at 10 rpm on an Economex rotarod fitted with a 3 cm-diameter rod (Columbus Instruments, Columbus, OH, USA), and the latency to fall (maximum 60 s) was measured to evaluate motor coordination and balance. Fixed speed rotarod assessment was performed at a constant speed of 10 rpm with a 300 s maximum time limit. After acclimation, all mice received training for 2 consecutive days. On the test day, the mice were tested in three consecutive trials of 1 min each, with 1 min rest between trials. The latency to fall during each of the three trials was averaged to give the overall time for each mouse.

*Measurement of grip strength in human subjects.* Grip strength was measured using a digital handheld dynamometer in a sitting position with elbows unsupported forming an angle of 90° (Lavisen, Hanam, Korea). Participants were asked to apply the maximum grip strength three times in the dominant hand. Between each measurement, at least 30 s of rest was allowed. Grip strength was defined as the maximally measured grip strength of the dominant hand.

*Transmission electron microscopy.* Gastrocnemius muscle samples from mice were fixed in 1% (wt/vol) glutaraldehyde at 4°C and then washed with 0.1 M cacodylate buffer, pH 7.2, at 4°C. Washed muscle tissues were fixed for 1 hour at 4°C with 1% (wt/vol) OsO4 in 0.1 M cacodylate buffer, pH 7.2, containing 0.1% (wt/vol) CaCl2. Muscle samples were dehydrated by graded series of ethanol and propylene oxide treatment, and then embedded in Embed-812 (Electron Microscopy Sciences). The resin blocks were then polymerized at 60°C for 48 hours. Tissues were sectioned with an EM UC6 ultramicrotome (Leica Microsystems, Vienna, Austria) and post-stained with 4% (wt/vol) uranyl acetate and citrate. Specimens were observed on a JEM ARM 1300S high-voltage electron microscope (JEOL, Japan).

### Legends to supplemental figures

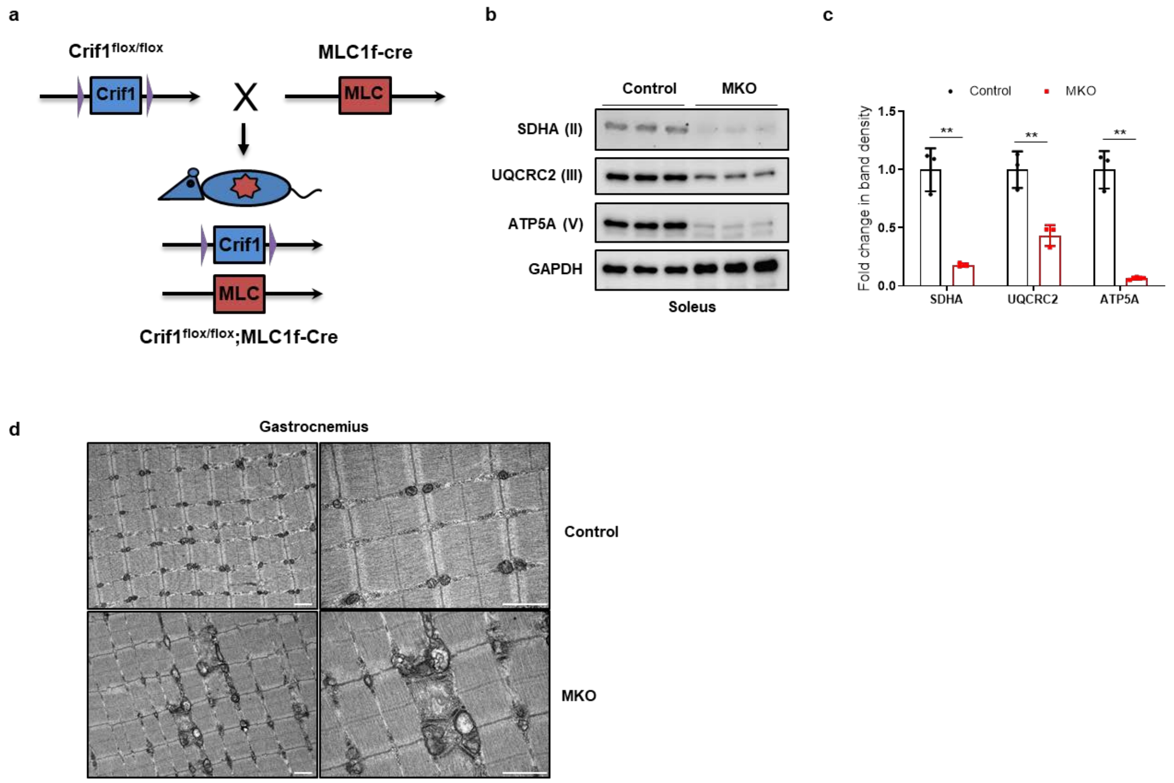


**Supplemental Figure 1. Generation of the skeletal muscle-specific mitochondrial oxidative phosphorylation (OxPhos) dysfunction mouse model.** (a) Strategy for generating MKO mice through disrupting *Crif1* in skeletal muscle using the MLC1f (myosin light chain 1f)-cre mice through Cre-loxP system. (b,c) Immunoblotting and band density measurement of OxPhos complex subunits in soleus muscle from 14-week-old control ($n$ = 3) and MKO ($n$ = 3) mice. (d) Mitochondrial morphology of controls and MKO mice, visualized by electron microscopy. Scale bars: 1 µm. Data are expressed as mean ± SEM. Statistical significance was analyzed by unpaired t-tests. **, $P < 0.01$ compared with the indicated group.

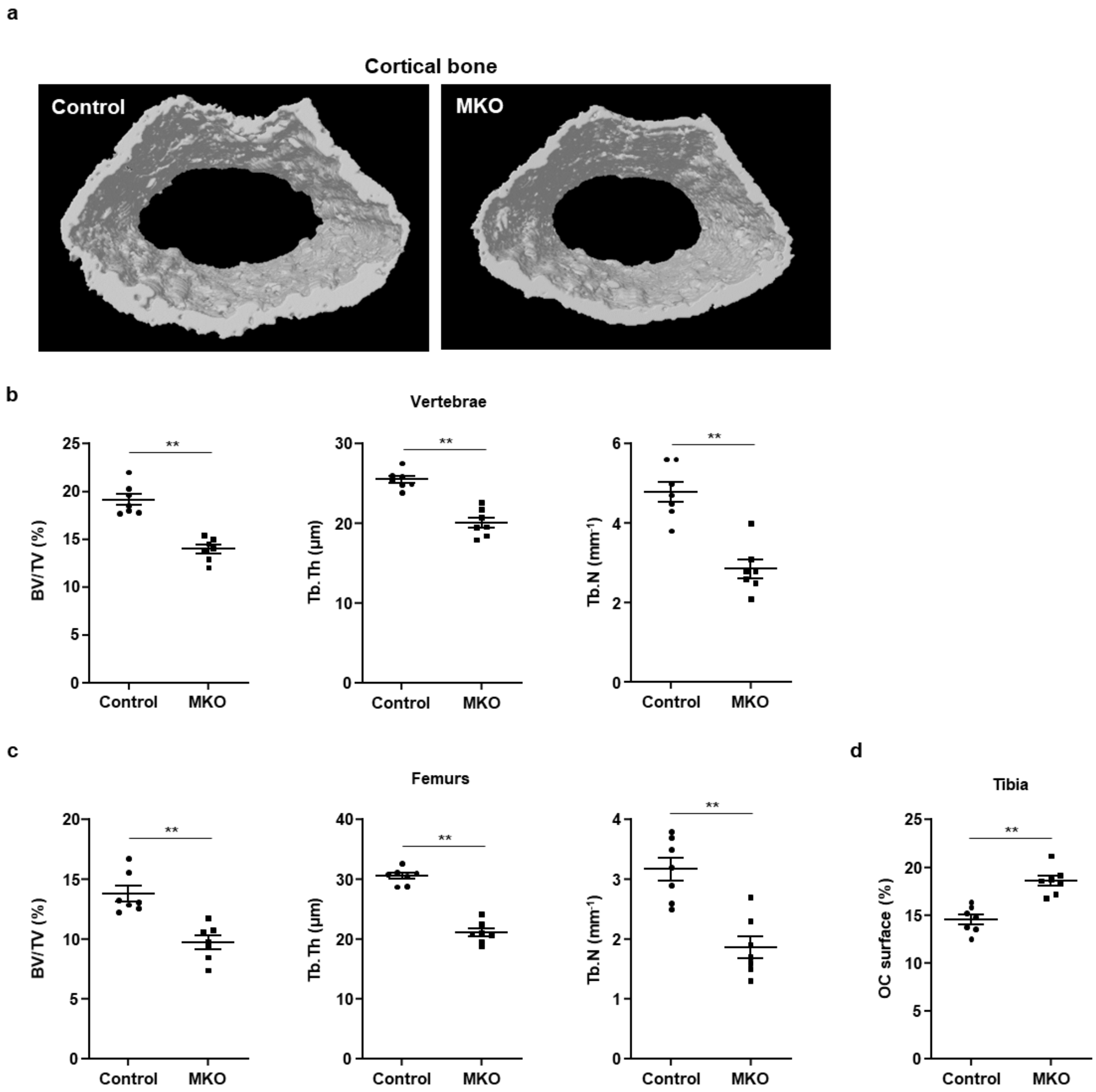


**Supplemental Figure 2. Micro-CT analysis of vertebrae and femurs of control and MKO mice at 14 weeks of age.** (a) The cortical area in femurs of control and MKO mice were measured by micro-CT. (b,c) Measurement of bone volume (BV)/total volume (TV) using von Kossa staining of vertebrae and femurs from 14-week-old control ($n$ = 5) and MKO ($n$ = 5) mice. (d) Osteoclast surface per bone surface (Oc.S/BS [%]) and number of osteoclasts per bone trabecular surface (N. Oc/Bpm [1 mm]) are indicated. Data are expressed as mean ± SEM. Statistical significance was analyzed by unpaired t-tests. *, $P < 0.05$ and **, $P < 0.01$ compared with the indicated group.

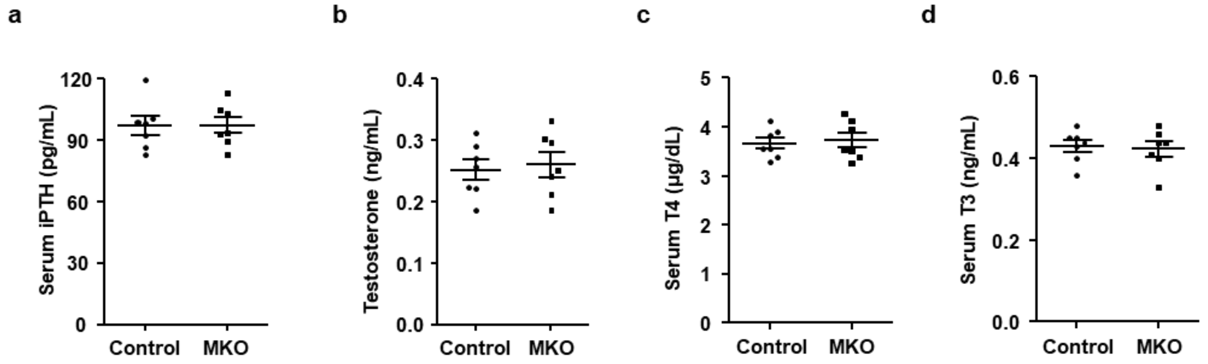


**Supplemental Figure 3. Measurement of serum levels of hormones affecting bone metabolism in control and MKO mice.** (a–d) Serum concentrations of intact parathyroid hormone (iPTH), testosterone, T4, and T3 in 14-week-old control and MKO mice ($n$ = 7/group). Data are expressed as mean ± SEM.

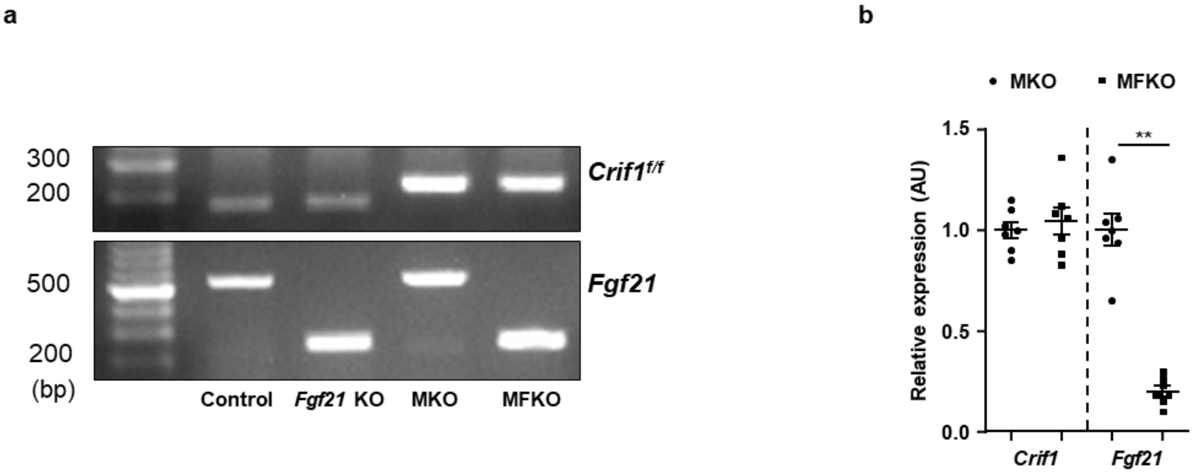


**Supplemental Figure 4. Generation of a *Fgf21*/*Crif1* double knockout mouse model.** (a) Strategy for generation of FGF21/CRF1 double knockout mice (MFKO) and genotype analysis using real-time PCR. (b) *Crif1* and *Fgf21* expression in the extensor digitorum longus muscle of MKO and MFKO mice at 6 weeks of age. Data are expressed as mean ± SEM. **, $P$ < 0.01 compared with the corresponding controls.

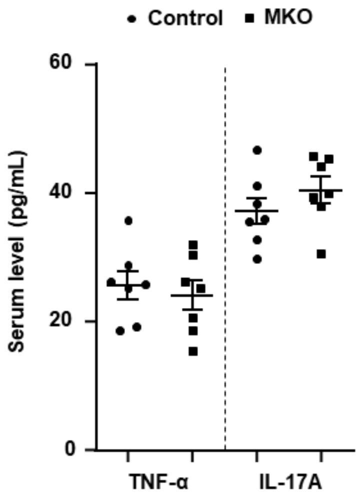


**Supplemental Figure 5. Measurement of serum levels of proinflammatory cytokines in 14-week-old control and MKO mice.** Serum TNF-α and IL-17A concentrations in control (*n* = 7) and MKO (*n* = 7) mice at 14 weeks of age. Data are expressed as mean ± SEM.

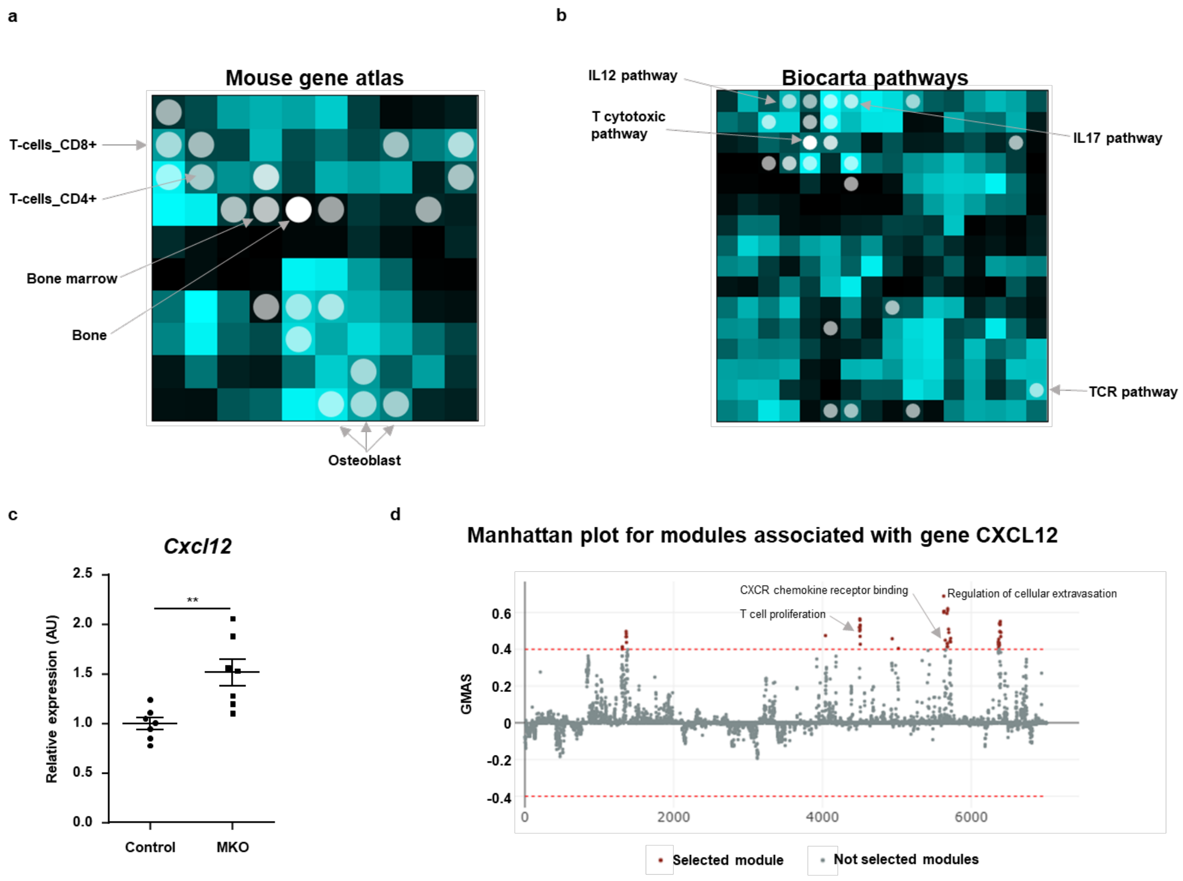


**Supplemental Figure 6. Analysis of RNA sequencing data from BM cells of control and MKO mice.** (a,b) The analysis was performed using Network2Canvas. Genes that were significantly upregulated in the BM cells of control and MKO mice were analyzed for gene-list enrichment, with gene set libraries created from level 4 of the MGI mouse phenotype ontology using Network2Canvas. (c) *Cxcl12* mRNA expression by BM cells from control and MKO mice at 14 weeks of age. (d) In the G-MAD analysis, CXCL12 associates with T-cell proliferation and CXCR chemokine receptor binding modules in mice. The threshold of significant gene-module association is indicated by the red dashed line. Modules are organized by module similarities. Known modules connected to CXCL12 are highlighted in red. Data are expressed as mean ± SEM. *, $P < 0.05$ and **, $P < 0.01$ compared with the corresponding controls.

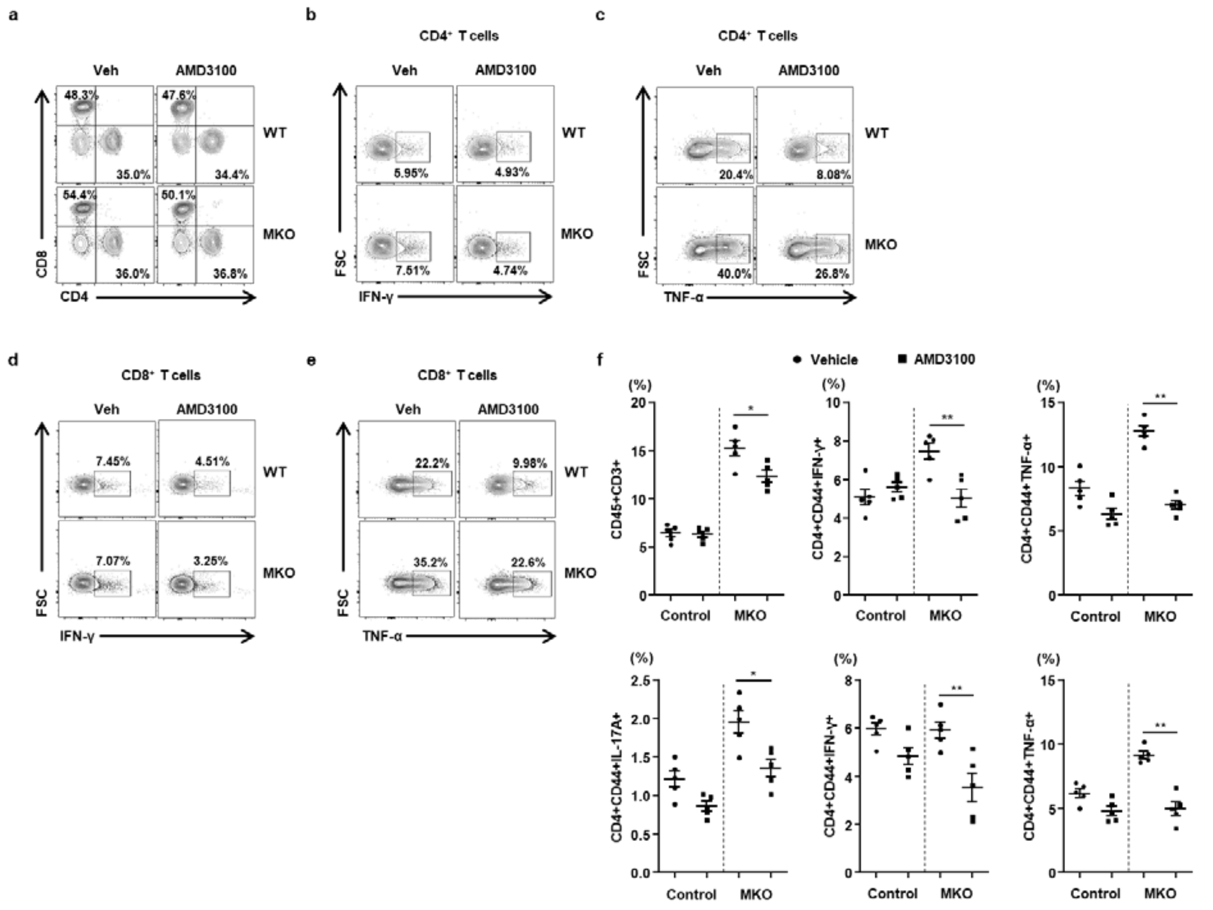


**Supplemental Figure 7. Flow cytometry analysis of BM cells from 12-week-old control and MKO mice treated with or without AMD3100.** (a) Populations of CD4+ and CD8+ T-cells from control and MKO mice treated with or without AMD3100 for 3 weeks. (b–e) IFN-γ- and TNF-α-producing CD4+ and CD8+ T-cells in the BM from control and MKO mice treated with or without AMD3100 for 3 weeks. (f) Statistical analysis of phenotypes defined by flow cytometry in the BM from control and MKO mice treated with or without AMD3100 for 3 weeks. Data are expressed as mean ± SEM. Statistical significance was analyzed by one-way ANOVA. *, $P < 0.05$ and **, $P < 0.01$ compared with the indicated group.

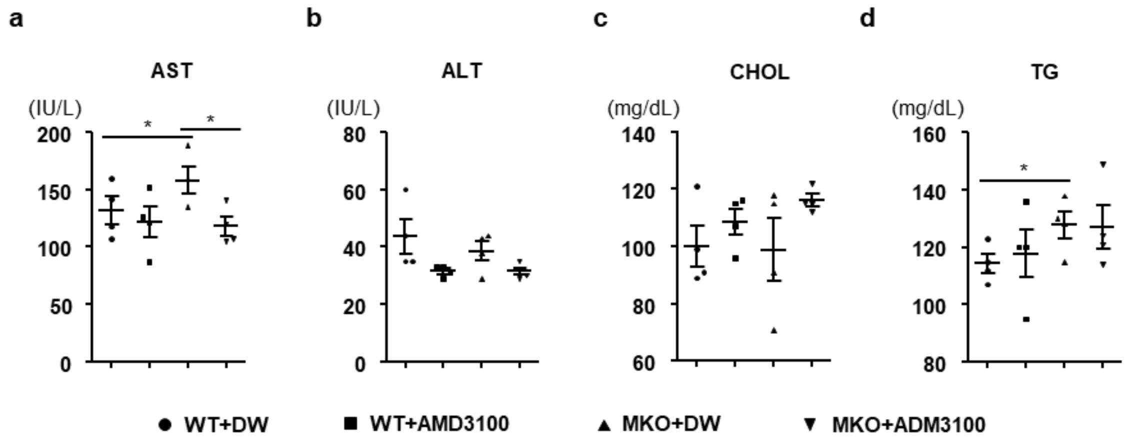


**Supplemental Figure 8. Measurement of serum markers of liver injury and lipid metabolism.** (a–d) Serum levels of aspartate transaminase (AST), alanine aminotransferase (ALT), total cholesterol (CHOL), and triglyceride (TG) from 12-week-old control and MKO mice treated with or without AMD3100. Data are expressed as mean ± SEM. Statistical significance was analyzed by one-way ANOVA. *, $P < 0.05$ and **, $P < 0.01$ compared with the indicated group.

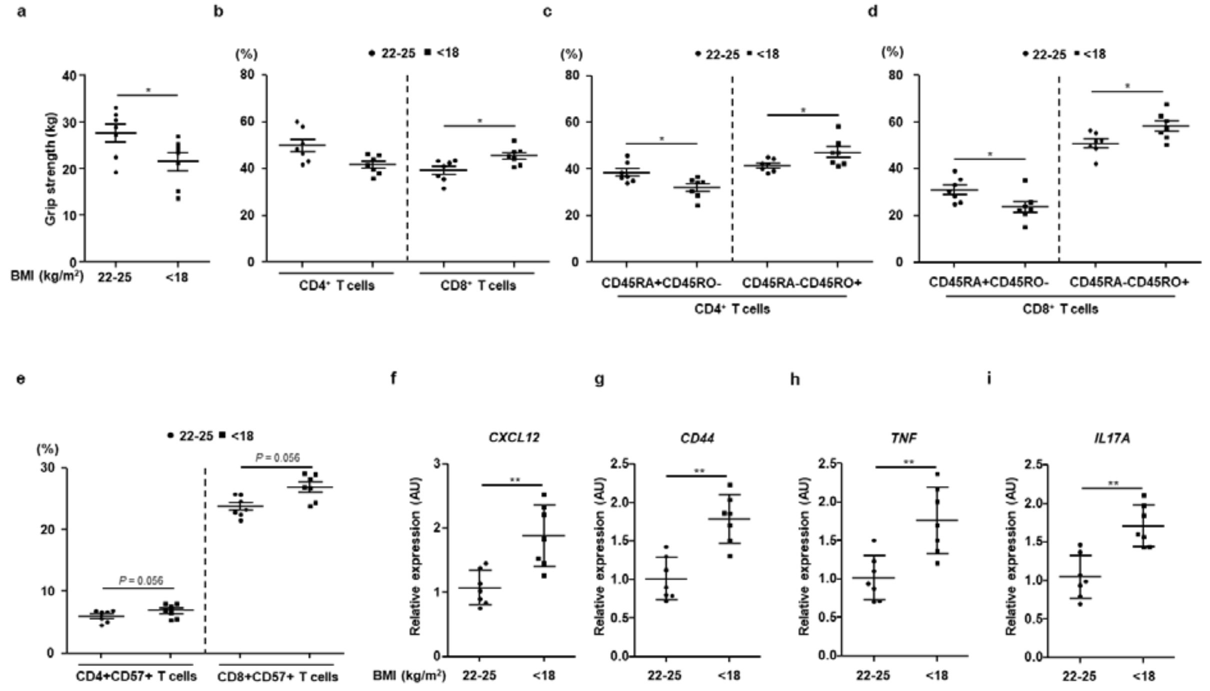


**Supplemental Figure 9. Grip strength and flow cytometry analysis of BM cells of patients with hip fracture.** (a) Measurement of grip strength of the hip fracture patients using a handheld dynamometer. (b) Populations of CD4+ and CD8+ T-cells from the hip fracture patients with lower (<18 $kg/m^2$) or normal (22–25 $kg/m^2$) body mass index (BMI). (c,d) Naïve (CD45RA+CD45RO-) and memory (CD45RA-CD45RO+) CD4+ and CD8+ T-cells in the BM from the hip fracture patients with lower (<18 $kg/m^2$) or normal (22–25 $kg/m^2$) BMI. (e) BM senescent (CD28-CD57+) CD4+ and CD8+ T-cells from hip fracture patients with lower (<18) or normal (22–25) BMI. (f–i) Transcript levels of *CXCL12*, *CD44*, *TNF*, and *IL17A* in the BM from hip fracture patients with lower (<18) or normal (22–25) BMI. Data are expressed as mean ± SEM. *, $P < 0.05$ and **, $P < 0.01$ compared with the corresponding controls.

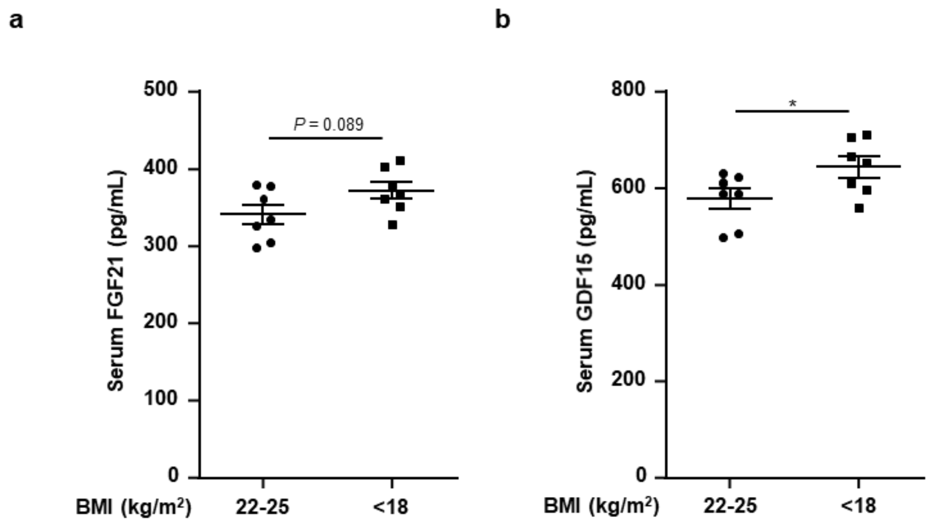


**Supplemental Figure 10. Serum levels of mitokines in patients with hip fracture.** (a,b) Serum FGF21 and GDF15 levels were measured in the hip fracture patients with lower (<18 kg/$m^2$) or normal (22–25 kg/$m^2$) BMI.

**Supplemental Table 1: Demographics and baseline characteristics of human subjects**

| BMI (kg/m$^2$) | No. | Gender | Age (years) | BMI | Grip strength (Kg) | Hip fracture site |
|---|---|---|---|---|---|---|
| 22–25 | 1 | M | 75 | 22.73 | 31.6 | Lt. intertrochanter |
| | 2 | F | 68 | 24.9 | 22.5 | Rt. neck |
| | 3 | M | 69 | 23.1 | 28.9 | Rt. neck |
| | 4 | F | 66 | 23.5 | 19.3 | Lt. intertrochanter |
| | 5 | M | 61 | 24.6 | 30.5 | Rt. intertrochanter |
| | 6 | M | 77 | 22.8 | 33.2 | Lt. neck |
| | 7 | M | 72 | 24.2 | 27.3 | Lt. neck |
| <18 | 1 | M | 78 | 17.6 | 25.3 | Rt. neck |
| | 2 | M | 61 | 17.1 | 23.6 | Rt. intertrochanter |
| | 3 | M | 80 | 17.5 | 21.2 | Rt. intertrochanter |
| | 4 | M | 66 | 16.8 | 26.9 | Lt. neck |
| | 5 | M | 65 | 17.3 | 24.4 | Rt. neck |
| | 6 | F | 72 | 17.6 | 15.2 | Lt. neck |
| | 7 | F | 70 | 16.5 | 13.6 | Lt. intertrochanter |

BMI, body mass index.

**Supplementary Table 2**. **Antibodies used for flow cytometry analysis**

| Antibody target/reagent | Fluorochrome | Clone | Supplier | Cat. No. |
|---|---|---|---|---|
| Fc block | | 2.4G2 | BD Bioscience | 553142 |
| Fixable viability dye | APC-Cy7 | | eBioscience | 65-0865 |
| hCD4 | Alexa Fluor 700 | RPA-T4 | eBioscience | 56-0049 |
| hCD8a | PE | RPA-T8 | eBioscience | 12-0088 |
| hCD8a | APC | RPA-T8 | eBioscience | 17-0088 |
| hCD3 | PerCP-Cy5.5 | SK7 | eBioscience | 46-0036 |
| hCD3 | PE-Cy7 | UCHT1 | eBioscience | 25-0038 |
| hCD57 | FITC | TB01 | eBioscience | 11-0577 |
| hCD28 | APC | CD28.2 | eBioscience | 17-0289 |
| hIFNG | PE-Cy7 | 4S.B3 | eBioscience | 25-7319 |
| hTNFA | APC | MAb11 | eBioscience | 17-7349 |
| hIL17A | APC | eBio64DEC17 | eBioscience | 17-7179 |
| mIL17A | PE | eBio17B7 | eBioscience | 12-7177 |
| mTNF-α | PerCP-eF710 | MP6-XT22 | eBioscience | 46-7321 |
| mTNF-α | APC | MP6-XT22 | eBioscience | 17-7321 |
| mIFNG | PE | XMG1.2 | eBioscience | 12-7311 |
| mIFNG | PE-Cy7 | XMG1.2 | eBioscience | 25-7311 |
| h/mCD44 | FITC | IM7 | eBioscience | 11-0441 |
| mNK1.1 | PE | PK136 | eBioscience | 12-5941 |
| mCD62L | APC | MEL-14 | eBioscience | 17-0621 |
| mCD4 | Per-cp-eF710 | GK1.5 | eBioscience | 46-0041 |
| hCD4 | FITC | RPA-T4 | eBioscience | 11-0049 |
| hCD3 | PE-eF610 | UCHT1 | eBioscience | 61-0038 |

| | | | | |
|---|---|---|---|---|
| hCD3 | SB436 | UCHT1 | eBioscience | 62-0038 |
| hCD8a | SB436 | RPA-T8 | eBioscience | 62-0088 |
| hTNFA | PE-Cy7 | Mab11 | eBioscience | 25-7349 |
| hIFNG | APC | 4S.B3 | eBioscience | 50-7319 |
| hCD45RA | FITC | HI100 | eBioscience | 11-0458 |
| hCD45RO | PE-Cy7 | UCHL1 | eBioscience | 25-0457 |
| hFOXP3 | APC | PCH101 | eBioscience | 17-4776 |
| h/mCD44 | SB436 | IM7 | eBioscience | 62-0441 |
| hTNFA | PE | Mab11 | eBioscience | 12-7349 |

**Supplemental Table 3. Primers used in real-time PCR (mouse)**

| Genes | Forward (5'–3') | Reverse (5'–3') |
|---|---|---|
| *Ppargc1a* | GAGCTACGGGGTCGCTTC | GGGACCCCAATCTCACCT |
| *Lonp1* | TGCGGAAACGCTAC AGGAC | GGAACAGAGCCCGGTGAAGG |
| *Clpp* | AAGCCTGTAGCCCACGTCGTA | AAGGTACAACCCATCGGCTGG |
| *Hspd1* | CCTCTCTCTAATCAGCCCTCTG | GAGGACCTGGGAGTAGATGAG |
| *Tid1* | GCCCATCCTCTGTGACTCAT | AGGCCACAGGTATTTTGTCG |
| *Chop* | TCCATCCAGTTGCCTTCTTG | TTCCACGATTTCCCAGAGAAC |
| *Atf4* | TCAGCCAGATGCAGTTAACGC | TCTGGACCCATTCCTTCTTGG |
| *Fgf21* | CAAAATCTCCAACCCATGCT | CACCACCAGGGTCTTCAAGT |
| *Tnf* | ATCGCAAACAAGCTGACCTG | AGATCCAGGTTTGAGGTGGG |
| *Rankl* | CCCTTGATGAAGAGGGATCA | ACTCCACAGGTGGGAACAAG |
| *Rorat* | TCCTCCAGGGATCCAACGA | GGCAGGCGGGAGGTCTT |
| *Rorgt* | CTGGGCTACACTGAGCACC | AAGTGGTCGTTGAGGGCAATG |
| *Il17a* | GCAAGAGATCCTGGTCCTGA | AGCATCTTCTCGACCCTGAA |
| *18s* | CTGGTTGATCCTGCCAGTAG | CGACCAAAGGAACCATAACT |

**Supplementary Table 4. Primers used in real-time PCR (human)**

| Genes | Forward (5'–3') | Reverse (5'–3') |
|---|---|---|
| *CXCL12* | CTGCCGCTTTGCAGGTGTA | CATTGTGGGCAAGGTGCTATT |
| *CD44* | ATTGTCCAGGCCAATACACATT | CCTCTCTACCTGCGTATCGTTTT |
| *TNF* | AAGGGGCAAAATGGTTCTTTCG | GCACCTGTATGTCCCCGAG |
| *IL17A* | TCGGTAACTGACTTGAATGTCCA | TCGCTTCCCTGTTTTAGCTGC |
| *ACTB* | CATGTACGTTGCTATCCAGGC | CTCCTTAATGTCACGCACGAT |
